# Structural basis for channel conduction in the pump-like channelrhodopsin ChRmine

**DOI:** 10.1101/2021.08.15.456392

**Authors:** Koichiro E. Kishi, Yoon Seok Kim, Masahiro Fukuda, Tsukasa Kusakizako, Elina Thadhani, Eamon F.X. Byrne, Joseph M. Paggi, Charu Ramakrishnan, Toshiki E. Matsui, Keitaro Yamashita, Takashi Nagata, Masae Konno, Peter Y. Wang, Masatoshi Inoue, Tyler Benster, Tomoko Uemura, Kehong Liu, Mikihiro Shibata, Norimichi Nomura, So Iwata, Osamu Nureki, Ron O. Dror, Keiichi Inoue, Karl Deisseroth, Hideaki E. Kato

## Abstract

ChRmine^1^, a recently-discovered bacteriorhodopsin-like cation-conducting channelrhodopsin^1, 2^, exhibits puzzling properties (unusually-large photocurrents, exceptional red-shift in action spectrum, and extreme light-sensitivity) that have opened up new opportunities in optogenetics^1, 3–5^. ChRmine and its homologs function as light-gated ion channels, but by primary sequence more closely resemble ion pump rhodopsins; the molecular mechanisms for passive channel conduction in this family of proteins, as well as the unusual properties of ChRmine itself, have remained mysterious. Here we present the cryo-electron microscopy structure of ChRmine at 2.0 Å resolution. The structure reveals striking architectural features never seen before in channelrhodopsins including trimeric assembly, a short transmembrane-helix 3 unwound in the middle of the membrane, a prominently-twisting extracellular-loop 1, remarkably-large intracellular cavities and extracellular vestibule, and an unprecedented hydrophilic pore that extends through the center of the trimer, separate from the three individual monomer pores. Electrophysiological, spectroscopic, and computational analyses provide insight into conduction and gating of light-gated channels with these distinct design features, and point the way toward structure-guided creation of novel channelrhodopsins for optogenetic applications in biology.

## Introduction

Light is one of the most important energy sources and environmental signals for living organisms, and motile organisms typically capture light using rhodopsin family proteins. Rhodopsins are largely classified into two groups: microbial (type 1) and animal (type 2), both consisting of a seven-transmembrane (7TM) protein termed the opsin, and a retinal chromophore covalently bound to the opsin via a conserved Schiff base; light absorption induces isomerization of the retinal and a series of subsequent photochemical reactions with conformational changes called the photocycle^6^. During the photocycle, microbial rhodopsins occupy a series of discrete and spectroscopically-distinguishable intermediate states (e.g. K, L, M, N, O), and ultimately transduce the photon by directly exerting biochemical functionality (diverse subfamily members include ion pumps, ion channels, sensors, histidine kinases, adenylyl/guanylyl cyclases, and phosphodiesterases^7, 8^). Heterologous expression of these proteins (especially of the ion-transporting channel- and pump-type rhodopsins) can be achieved by targeting the opsin genes to specific cell types, which (when applied along with precise light-delivery technology) enables high-speed control of the membrane potential of specific cells in behaving organisms with light. This experimental approach, called optogenetics, has been broadly applied to provide or inhibit activity patterns in specific cells, enabling targeted causal study of cellular activity in living systems^9, 10^.

Both channel- and pump-type rhodopsins are well-established in optogenetics, and cation channelrhodopsins (CCRs) are typically used for activation of target cells (by depolarizing the cellular membrane potential^11^). The initial biophysical description of a CCR (*Cr*ChR1 from the chlorophyte *Chlamydomonas reinhardtii*) in 2002^12^ was followed by characterization of many CCR variants that were discovered in nature and/or redesigned as optogenetic tools with new types of functionality spanning ion selectivity, photocurrent amplitude, absorption spectrum, light sensitivity, and on/off-kinetics^9, 11^. Natural CCRs include *Cr*ChR2 (channelrhodopsin-2 from *Chlamydomonas reinhardtii*)^12^, VChR1 (channelrhodopsin-1 from *Volvox carteri*)^13^, *Co*ChR (channelrhodopsin from *Chloromonas oogama*)^14^, Chronos (channelrhodopsin from *Stigeoclonium helveticum*)^14^, and Chrimson (channelrhodopsin from *Chlamydomonas noctigama*)^14^; these were initially described chiefly from chlorophyte algae, but identification of new channelrhodopsins from species beyond chlorophytes further expanded the diversity of optogenetic tools. In 2016-7, a distinct subfamily of microbial rhodopsins was reported from cryptophyte *Guillardia theta*^15, 16^. These cryptophyte rhodopsins were biophysically found to be light-gated cation channels, but are encoded by genes more homologous to those of archaeal ion-pumps (such as *Halobacterium salinarum* bacteriorhodopsin or *Hs*BR) than canonical chlorophyte CCRs. Moreover, unlike chlorophyte CCRs, these cryptophyte CCRs share three amino acids on TM3 that are crucial for outward proton pumping (the DTD motif^17^; D85, T89, and D96 in *Hs*BR) and accordingly have been termed bacteriorhodopsin-like cation channelrhodopsins or BCCRs^2^ (Extended Data Fig. 1). ChRmine (a BCCR discovered through a structure-guided genome mining approach^1^) exhibits extremely high photocurrents and light sensitivity, as well as a markedly red-shifted action spectrum; these properties have significantly reshaped the application domain of optogenetics, including in behaving mammals, from all-optical interrogation of hundreds of individually-specified single neurons^1^ to fully non-invasive transcranial control of deep brain neural circuitry^3^.

Despite the importance of these properties, both for optogenetic applications and for exploring microbial opsin diversity and evolution, there is no structure available for ChRmine or any BCCR. Such high-resolution structural information would be of enormous value, not only to enhance our fundamental understanding of the diverse structure-function relationships among pump- and channel-type rhodopsins, but also to provide a framework for creation of next-generation optogenetic tools (a process which, for example, previously led to creation of the initial anion-conducting channelrhodopsins^18–21^). Here we present the cryo-electron microscopy (cryo-EM) structure of ChRmine at 2.0 Å resolution, which reveals (unlike any other known channelrhodopsin) a BR-like trimer configuration and an unusually short TM3 unwound from the middle of the membrane. The long extracellular loop-1 (ECL1) arising from TM3 unwinding contributes to creation of deformed structure in the Schiff base region, a large extracellular cavity, and ion selectivity filter-like narrowing at the center of the trimer interface. Electrophysiological and computational analyses suggest that, in addition to the canonical ion-conducting pathway within each monomer, a hydrophilic pore within the trimer interface can operate as an auxiliary ion-conducting pathway.

## Structural determination

Our initial efforts to crystallize ChRmine yielded inadequately-diffracting crystals; we therefore turned to single-particle cryo-EM (Extended Data Fig. 2). ChRmine is a small (∼35kDa) and compact membrane protein without a large extracellular or intracellular domain, so the particles picked up from an initial cryo-EM dataset were not well-aligned and failed to yield three-dimensional reconstructions (Extended Data Fig. 2a-d). We therefore generated antibody fragments against ChRmine and obtained one conformation-specific antibody Fab02 (Methods; Extended Data Fig. 3a and b). To assess the effect of Fab02 on ChRmine, we spectroscopically analyzed the ChRmine-Fab02 complex alongside ChRmine and confirmed that Fab02 binding did not affect the photocycle; both ChRmine and ChRmine-Fab02 complexes exhibited K, L_1_, L_2_, M_1,_ and M_2_ intermediates with similar lifetimes (Extended Data Fig. 3c-f, 4 and Supplementary Table 1). Using this Fab fragment, the structure of the ChRmine-Fab02 complex in the dark state was determined at an overall resolution of 2.0 Å (Extended Data Fig. 2e-j, 3g,h, and Supplementary Table 2). The density was of excellent quality and allowed for accurate modelling of ChRmine continuously from residues 10 to 279 (Extended Data Fig. 5a-d; Methods). The density also clearly resolved several lipids, water molecules, and the all*-trans*-retinal (with the specific retinal conformer confirmed by high-performance liquid chromatography; Extended Data Fig. 5a-d and 6); signals of putative hydrogen atoms were even observed in the difference (*F_o_-F_c_*) map (Extended Data Fig. 5e and f)^22^. As far as we know, this initial BCCR structure is also one of the highest-resolution structures of any channelrhodopsin (indeed, the first rhodopsin for which some hydrogen atoms can be visualized).

## Overall structure and comparison with *Hs*BR and C1C2

The cryo-EM density map first revealed that the quaternary structure of ChRmine is strikingly different from that of other structure-resolved channelrhodopsins^18^ (Fig. 1a-c). Instead of the classical channelrhodopsin dimer, ChRmine rather forms a trimer, and TM2 interacts with TM4 of adjacent protomers as observed in archaeal ion-pumping rhodopsins including *Hs*BR^23^ and *Hs*HR^24^ (Extended Data Fig. 7a-c). To confirm this result under more native conditions, we next reconstituted ChRmine in a lipid bilayer and performed high-speed atomic force microscopy (HS-AFM) analysis. The HS-AFM imaging clearly revealed trimeric structure as well, indicating that ChRmine assembles as a trimer not only in detergent micelles but also in a physiological bilayer environment (Extended Data Fig. 7d).

**Fig. 1.**
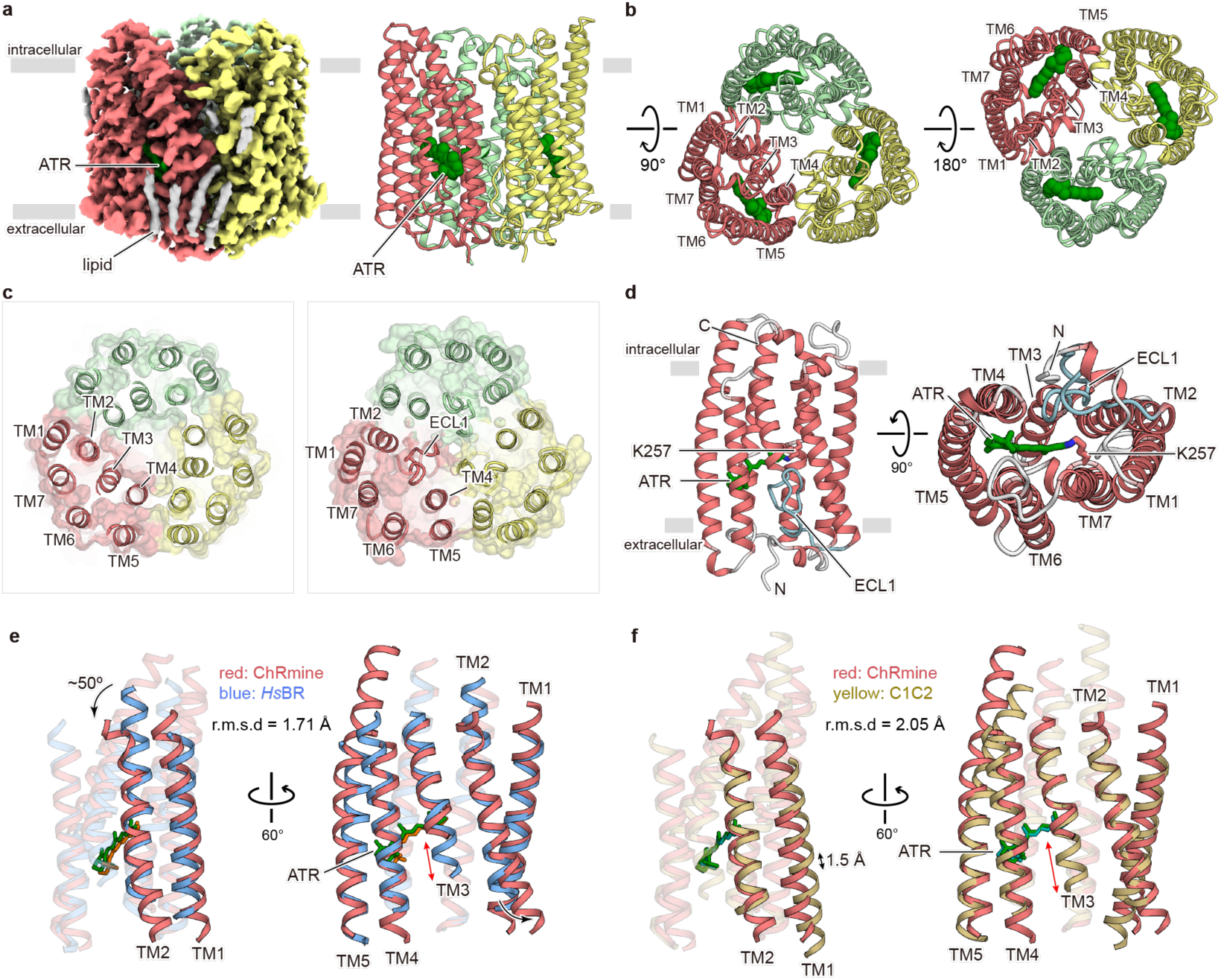
Cryo-EM structure of ChRmine and comparison with *Hs*BR and C1C2. **a**, Cryo-EM density map (left) and ribbon representation (right) of ChRmine homotrimer, coloured by protomer (red, yellow, and light green), ATR (green), and lipid (grey). **b**, Ribbon representation of ChRmine homotrimer, viewed from the intracellular side (left) and extracellular side (right). **c**, Cross sectional view of the intracellular side (left) and extracellular side (right) of ChRmine homotrimer. Semi-transparent molecular surface is overlaid. **d**, ChRmine monomer, viewed parallel to the membrane (left), and from the extracellular side (right). ATR, depicted by the stick model, is coloured green, and ECL1 is coloured cyan. **e**, **f**, ChRmine (red) superimposed onto *Hs*BR (blue) (**e**) and C1C2 (yellow) (**f**) from two angles. Single-head black arrows, double-head black arrows, and double-head red arrows highlight TM2 bending, TM1 shifting, and TM3 shortening, respectively.

The monomeric structure of ChRmine consists of an extracellular N-terminal domain (residues 10-26), an intracellular C-terminal domain (residues 271-279), and 7 TM domains (within residues 27-270), connected by three intracellular loops (ICL1 to ICL3) and three extracellular loops (ECL1 to ECL3) (Fig. 1d). TM1–7 adopt a canonical rhodopsin-like topology with a covalently-linked retinal at K257 on TM7, but TM3 markedly diverges from the classical framework, exhibiting an unwound configuration in the middle of the transmembrane region, leading to a long twisting ECL1 (residues 95–115) and a resulting C-shaped structure that is stabilized by an extensive hydrogen-bonding network (Extended Data Fig. 5d).

To further explore how ChRmine can be structurally similar to ion-pumping rhodopsins and yet function as a cation channel, we next closely analyzed the individual-monomer structural features of ChRmine in comparison with representatives of an archaeal ion-pumping rhodopsin (*Hs*BR), and a chlorophyte CCR (C1C2, the chimeric channelrhodopsin derived from *Cr*ChR1 and *Cr*ChR2). Consistent with the sequence similarity (Extended Data Fig. 1a and b), ChRmine can be better superimposed onto *Hs*BR; the root-mean-square deviation (r.m.s.d) values of ChRmine vs. *Hs*BR and C1C2 were measured to be 1.71 Å and 2.05 Å, respectively (Fig. 1e and f). Notably, while previous structural work had revealed that TM1, 2, 3, and 7 of CCRs form the ion-conducting pathway within each monomer and that positioning of TM1 and 2 structurally distinguishes channelrhodopsins from pump-type rhodopsins^18^, TM1 of ChRmine is positioned more similarly to that of *Hs*BR, and is shifted in its entirety by 1.5 Å in ChRmine relative to C1C2 (Fig. 1f). The overall positioning (and the central region) of TM2 is also similar between ChRmine and *Hs*BR, with the exception that both the intracellular and extracellular regions of TM2 are tilted outward in ChRmine (Fig. 1e); these features in TM2 enlarge the cavity within the monomer and may allow ChRmine to function as a cation channel (addressed in detail later).

## The Schiff base region

In all microbial rhodopsins, the retinal is covalently bound to a TM7 lysine residue, forming the protonated Schiff base; the positive charge of the Schiff base proton is stabilized by 1-2 carboxylates located at the extracellular side (Extended Data Fig. 8a). After photon absorption, the proton is transferred to the carboxylate; this proton movement is a critical step in the function of most ion-transporting rhodopsins^6^. The carboxylate(s) stabilizing the positive charge and receiving the proton (forming the M intermediate) are historically termed the “Schiff base counterion(s)” and “proton acceptor”, respectively^6^. To obtain structural insight into ChRmine channel gating mechanisms and dynamics, we next focused on the counterion and proton acceptor.

Surprisingly, the Schiff base region of ChRmine is remarkably different from that of both pumping rhodopsins and other channelrhodopsins (*Hs*BR and C1C2; Fig. 2a), despite the fact that the primary sequence, the oligomerization number, and the monomeric overall structure of ChRmine are similar to *Hs*BR (Extended Data Fig.1, 7 and Fig 1). In *Hs*BR, the protonated Schiff base nitrogen hydrogen-bonds with a water molecule between the Schiff base counterions, D85 and D212. D212 is fixed by hydrogen bonds with Y57 and Y185 on TM2 and 6, respectively, while D85, which works as the proton acceptor from the Schiff base in the M intermediate^25^, interacts with R82 via water molecules (Fig. 2a middle). In C1C2, D85, Y57, and Y185 in *Hs*BR are replaced by E162, F133, and F265, respectively, and the Schiff base nitrogen hydrogen-bonds with D292, which no longer interacts with F133 and F265 (Fig. 2a right). D292 works as the proton acceptor in the M intermediate^18, 26^, and E162 is dispensable for channel function^18, 27^. In contrast, in ChRmine, while the residues corresponding to Y57, R82, D85, and D212 in *Hs*BR are conserved (Y85, R112, D115, and D253 in ChRmine), R112 and D115 are displaced far from the Schiff base due to the unwinding of TM3 (Fig. 2a left); the distance from the Schiff base to R112 and D115 is 13.7Å and 6.8Å (11.2 Å and 7.8 Å for Cα), respectively, and such long distances have never been observed in other microbial rhodopsins with solved structures (Extended Data Fig. 8b-d).

**Fig. 2.**
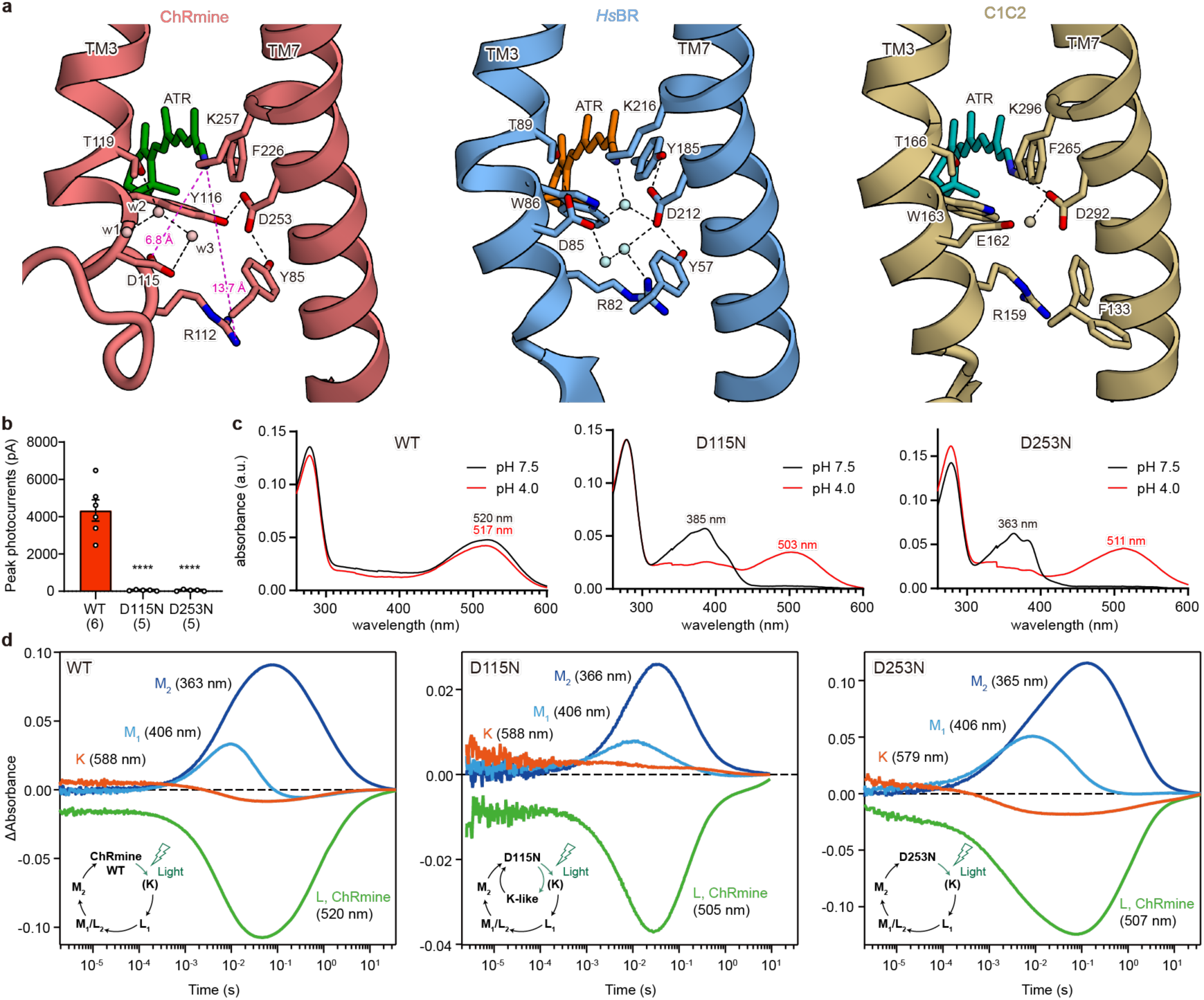
The Schiff base region. **a**, The Schiff base regions of ChRmine (left), *Hs*BR (middle), and C1C2 (right). Spheres and black dashed lines represent water molecules and hydrogen bonds, respectively. **b**, Photocurrent amplitudes of wild-type (WT) ChRmine and two mutants. Both mutants abolish channel activity. Data are mean ± s.e.m; one-way ANOVA followed by Dunnett’s test. \*\*\*\**P* < 0.0001. Sample size (number of cells) indicated in parentheses. **c**, Absorption spectra of ChRmine WT (left), D115N (middle), and D253N (right) at pH 7.5 (black) and pH 4.0 (red). Note that the Schiff base is deprotonated at pH 7.5 and re-protonated at pH 4.0 in D115N and D253N mutants, suggesting that both deprotonated D115 and D253 are required to stabilize the Schiff base proton at pH 7.5. **d**, Time-series traces of absorption changes for ChRmine WT (left), D115N (middle), and D253N (right) at 363-366 (blue), 406 (cyan), 505-520 (green), 579-588 nm (red) probe wavelengths. Inset shows photocycle schemes as determined by flash photolysis. Only the D115N mutant exhibits an additional K-like intermediate.

Three water molecules, w1, w2, and w3, occupy the space between the Schiff base and D115 created by the unwinding of TM3. Notably, w2 and w3 are well superposed onto the carboxyl oxygens of D85 in *Hs*BR, suggesting that these water molecules structurally mimic D85 (Extended Data Fig. 8b) and participate in the counterion complex with D115 and D253. In addition to these structural changes in the TM3 region, the substitutions of Y185 (in *Hs*BR) to F226, and W86 (in *Hs*BR) to Y116, also rearrange the structure in the TM7 region of ChRmine; D253 switches the hydrogen bond from F226 to Y116, and unlike *Hs*BR, D253 in ChRmine is fixed by two tyrosines (Y85 and Y116) on TM2 and 3 (Fig. 2a left).

To explore the function of the counterion and proton acceptor candidates, D115 and D253, we measured photocurrent amplitudes of wild-type (WT), D115N, and D253N ChRmine in human embryonic kidney 293 (HEK293) cells. Whereas D115N and D253N mutants exhibited robust expression and membrane localization of the rhodopsin protein itself, both mutants completely abolished photocurrents (Fig. 2b and Extended Data Fig. 9). Spectroscopic analysis revealed that these mutants have strikingly blue-shifted absorption spectra (*λ*_max_ at pH 7.5 shifting from 520 nm (WT) to 385 nm (D115N) and 363 nm (D253N), respectively), consistent with the loss of function arising from baseline deprotonation of the Schiff base (all at physiological pH; Fig. 2c). This is the pattern expected if D115 and D253 work as the Schiff base counterions that must be deprotonated at baseline under physiological pH for essential stabilization of the positive charge of the protonated Schiff base. In other words, both D115 and D253 work as the Schiff base counterions.

Next, to identify which carboxylate works as a primary proton acceptor in the M intermediate, we performed flash photolysis of D115N and D253N mutants (Fig. 2d). Because rhodopsins with the Schiff base deprotonated cannot respond to light, the photocycles of D115N and D253N mutants were measured under acidic conditions (pH 4.0) to re-protonate the Schiff base (Fig. 2c). Spectroscopic analysis revealed that both mutants have similar photo-intermediates compared to WT, but only the D115N mutant shows additional accumulation of a K-like intermediate (long-lived up to one second) with lower accumulation of M intermediates (Fig. 2d). Indeed, the decay time of the M_2_ intermediate was significantly faster in D115N (τ_M2_ = 190 ± 40 ms) compared to WT (τ_M2_ = 1.09 ± 0.06 s) (Supplementary Table 1), consistent with D115 operating as a primary proton acceptor. While D253 is located closer to the Schiff base than D115, the 253 aspartate strongly interacts with Y85 and Y116, which would make it difficult for D253 to receive the proton from the Schiff base. D212 of *Hs*BR similarly interacts with two tyrosine residues (Y57 and Y185) and also does not act as the proton acceptor. D115 is located further from the Schiff base but with several water molecules positioned in between; water rearrangements would allow the proton would be transferred from the Schiff base to D115 in the M intermediate.

## Ion-conducting pore within the monomer

To explore the location and shape of the ion-conducting pathway, we first analyzed the configuration of cavities within the ChRmine monomer. By comparison with C1C2 and *Hs*BR, ChRmine displays markedly larger intracellular and extracellular cavities (Fig. 3a). As in C1C2, both cavities are mainly formed by TM1, 2, 3, and 7, and are occluded by intracellular and central constriction sites (ICS and CCS); however, multiple key differences in the pore pathways of ChRmine and C1C2 were noted. First, while computed electrostatic surface potentials for both ChRmine and C1C2 reveal electronegative pores (Extended Data Fig. 10a-c), the distributions of negatively-charged residues were found to be remarkably different. In C1C2 and several other chlorophyte CCRs, five conserved glutamates (E121, E122, E129, E136, and E140 in C1C2) cooperatively create the electronegative surface potential along the pore (Extended Data Fig. 10c and d), but four of these five residues are substituted with neutral or basic residues in ChRmine (Extended Data Fig. 1a and 10a). Instead, ChRmine displays a distinct set of carboxylates including E50, E70, D100, D126, E154, E158, D242, E246, and D272, to create cavities suitable for anion exclusion and cation selectivity^19, 21, 28^ (Extended Data Fig. 10b).

**Fig. 3.**
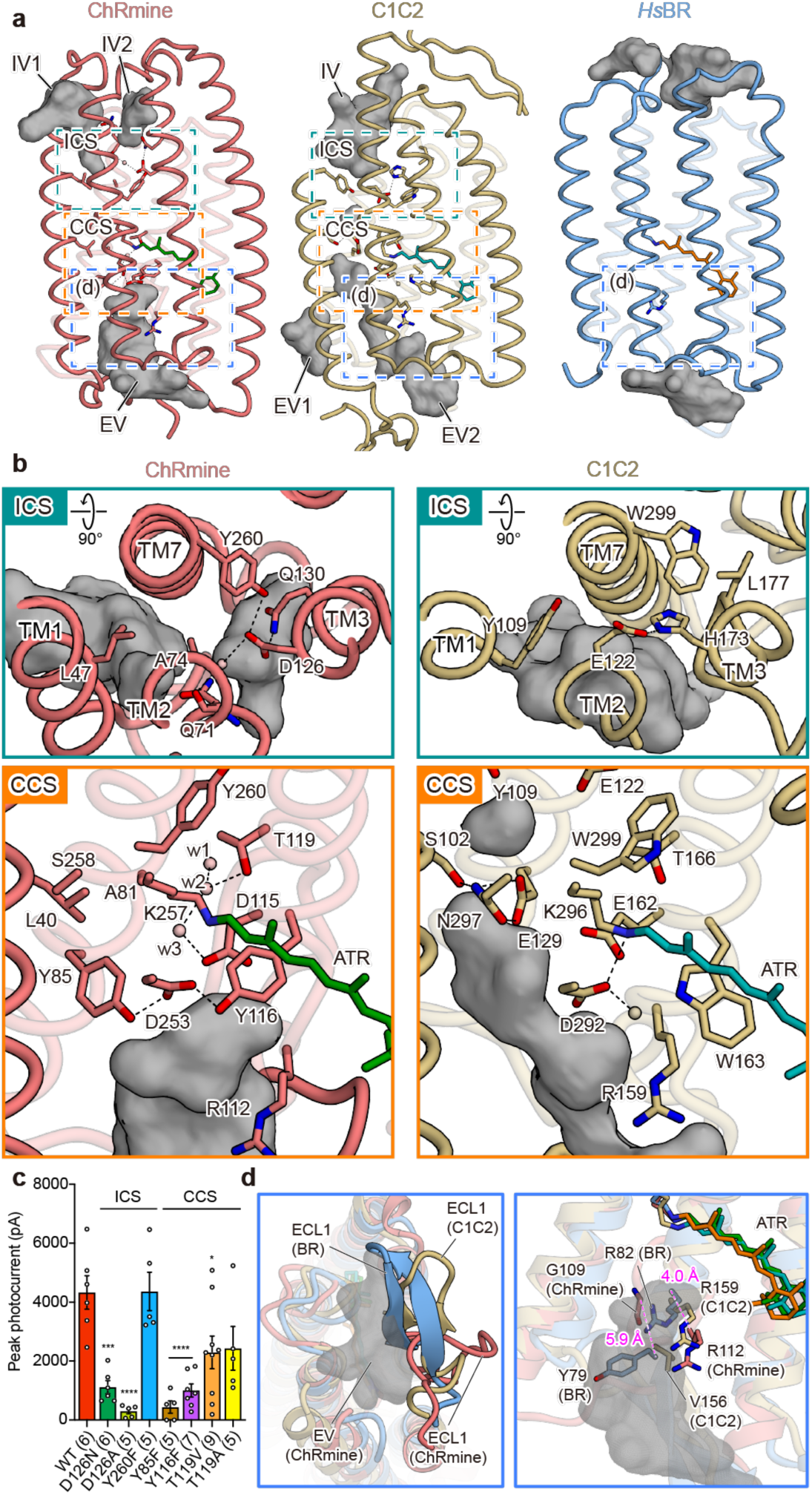
Ion-conducting pathway within the monomer. **a**, The extracellular and intracellular vestibules (EV and IV, respectively) of ChRmine (left), C1C2 (middle), and *Hs*BR (right) within the monomer. Vestibules are coloured in grey. The intracellular and central constriction sites (ICS and CCS) are indicated by dashed boxes (green and orange, respectively). Note that only ChRmine and C1C2 have ion-conducting hydrophilic pores within the monomer. **b**, ICS (top) and CCS (bottom) of ChRmine (left) and C1C2 (right). **c**, Effects of mutations in constriction sites. Data are mean ± s.e.m; one-way ANOVA followed by Dunnett’s test. * P < 0.05, \*\*\*\**P* < 0.0001. Sample size (number of cells) indicated in parentheses. **d**, Magnified views of the blue-boxed region in **a**. Comparison of the overall structure (left) and key residues (right) of ECL1 between ChRmine (red), C1C2 (yellow), and *Hs*BR (blue). The extracellular cavity of ChRmine is depicted as grey mesh. ECL1 of ChRmine adopts the different conformation shown (left), while in *Hs*BR Y79 and R82 efficiently occlude the cavity.

Second, ChRmine exhibits two intracellular vestibules (IV) with distinct electrostatic potentials (Fig. 3a left and Extended Data Fig. 10a). Notably, the position of ChRmine IV1 is more similar to the IV of the cation channelrhodopsin C1C2, and the position of ChRmine IV2 is more similar to the IV of the anion channelrhodopsin *Gt*ACR1^19, 25^ (Fig. 3a and Extended Data Fig. 10a and d). The corresponding electrostatic surface potentials favor a role for IV2 as a cation conducting pore in the open state (IV1 and IV2 could further connect to create a larger intracellular cavity in the open state, as suggested for extracellular vestibules of other channelrhodopsins^29, 30^).

Third, the ICS architecture of ChRmine and C1C2 are different. In C1C2, the ICS is mainly formed by Y109, E122, and H173 (E122 and H173 are hydrogen-bonded to each other). In ChRmine, the corresponding residues are L47, A74, and D126, respectively, which participate in the formation of the ICS, but D126 forms a more extensive hydrogen-bonding network with Q71, Q130, Y260, and a water molecule (Fig. 3b). While mutation of Y260 does not compromise channel activity, D126 mutants show severely decreased photocurrents, suggesting that the effect of loss of a single hydrogen bond at the ICS is minimal, while even a small change to D126 as the hub of the hydrogen-bonding network significantly affects channel activity (Fig. 3c and Extended Data Fig. 9). Fourth, the size and path of the extracellular cavities significantly differ between ChRmine and C1C2. C1C2 has two extracellular vestibules (EV1 and EV2), but ChRmine lacks the vestibule corresponding to EV1, while the volume of ChRmine’s sole EV is significantly expanded (due in large part to TM3 unwinding; Fig. 3a). In addition, the EV2 of C1C2 is well-separated from the Schiff base and terminates at the CCS formed by S102, E129, and N297; in contrast, the EV of ChRmine extends prominently to the Schiff base region (Fig. 3b), and ChRmine’s three residues corresponding to the CCS of C1C2 (L40, A81, S258) do not form a constriction. Instead, the extensive hydrogen-bonding network formed by the counterion complexes (including D115, D253, Y85, Y116, T119, and structured water molecules) occlude the pore and define the ChRmine CCS; the importance of this hydrogen-bonding network is supported by the loss-of-function electrophysiological properties of Y85F, Y116F, and T119V mutant photocurrents (Fig. 3c).

While ChRmine resembles *Hs*BR in primary sequence, overall arrangement of the secondary structural elements of the monomer, and quaternary structure of the trimer (Fig. 1 and Extended Data Figs. 1 and 7), the size and shape of the cavities within the monomer clearly show higher similarity to those of C1C2, consistent with the cation channel functionality of ChRmine (Fig. 3a). We next sought to understand which structural elements contribute to formation of these large cavities that comprise much of the channel pore in ChRmine, by comparing ChRmine and *Hs*BR in more detail. At least two notable features contribute to this formation of the pore structure. First, as described above, both ends of TM2 are tilted outward in ChRmine; the cytoplasmic end of TM2 is particularly tilted, by about 50 degrees, which significantly enlarges the intracellular cavity (Fig.1e and Extended Data Fig. 10e). Furthermore, numerous hydrophilic residues (including S54, E70, Q71, D126, Q130, R268, and D272) face into the pore, which together with the structural waters creates an environment suitable for water and ion conduction. In contrast, in *Hs*BR TM2 is straight through the end, and 6 of the above 7 hydrophilic residues are replaced by hydrophobic residues, which are tightly packed with no water-accessible cavity (Fig. 1e, Fig. 3a right, and Extended Data Fig. 10f).

In a second major channel-enabling feature, the unwinding of TM3 and resulting long ECL1 contribute to creation of a large extracellular cavity in ChRmine. The helical structure of extracellular TM3 is unfolded beginning at Y116, and the C-shaped structure of ECL1 protrudes to the center of the trimer interface. This is in contrast to the ECL1 of *Hs*BR, which forms a β-sheet and is located in a position that half-occludes the extracellular pore (Fig. 3d left). In addition to the overall position of ECL1, R82 on TM3 and Y79 on ECL1 protrudes into and occludes the extracellular cavity in *Hs*BR (Fig. 3d right). However, Y79 is replaced by G109 in ChRmine, and because of the unfolding of TM3, R112 (R82 in *Hs*BR) and G109 are displaced by 4.0 Å and 5.9 Å, respectively, from the corresponding residues of *Hs*BR. As a result, these residues do not block the cavity in ChRmine (Fig. 3d right).

Notably, the ECL1 of C1C2 also forms a β-sheet structure like *Hs*BR and moderately narrows the entrance of the pore– one of the reasons that the extracellular cavity of C1C2 is smaller than that of ChRmine (Fig. 3d left). Moreover, in C1C2, while the residues corresponding to R82 and Y79 of *Hs*BR are located in similar positions, Y79 is replaced by V156, and R159 (R82 in *Hs*BR) adopts the conformation similar to that of R112 in ChRmine, facing towards the extracellular solvent rather than running parallel to the membrane (Fig. 3d right). The outward-facing Arg conformation observed in ChRmine and C1C2 is conserved in other structure-known channel-type rhodopsins including *Cr*ChR2, C1Chrimson, and *Gt*ACR1, and the parallel Arg conformation observed in *Hs*BR is also generally conserved in other pump-type rhodopsins such as halorhodopsins (inward Cl^-^ pump-type), KR2 (outward Na^+^ pump-type), and schizorhodopsin (inward H^+^ pump-type) (Extended Data Fig. 8d). These results suggest that, as well as the overall size of the cavity within the monomer, the outward-facing arginine conformation in the dark state is a key structural element defining function of ion-transporting rhodopsins, consistent with previous studies showing that mutating this arginine converts ion-pump to ion-channel rhodopsins^31, 32^ (Supplementary discussion).

## Hydrophilic pore at the trimer interface

Like *Hs*BR, ChRmine forms a trimer; here we find that ChRmine has an unprecedented additional pore at the trimer interface (Fig. 4a). The corresponding region in *Hs*BR is hydrophobic and filled with several lipid molecules, but for ChRmine this region is hydrophilic and negatively charged (Extended Data Fig. 11a-d). The chalice-shaped pore is formed by TM2-4 and ECL1 of the protomer, and the narrowest region is created by ECL1; the main-chain carbonyl oxygens of F104 and I106 face toward the pore center and form the constriction (Fig. 4b and c). The C-shaped architecture of each ECL1 (Fig. 4c left) and the trimeric assembly of the carbonyl oxygen (Fig. 4c right) resemble the selectivity filters of the tetrameric potassium channel (e.g. KcsA and MthK) and the trimeric cation channel (e.g. ASIC and ENaC), respectively (Extended Data Fig. 11e, f), suggesting that the central pore functions as an auxiliary ion-conducting channel.

**Fig. 4.**
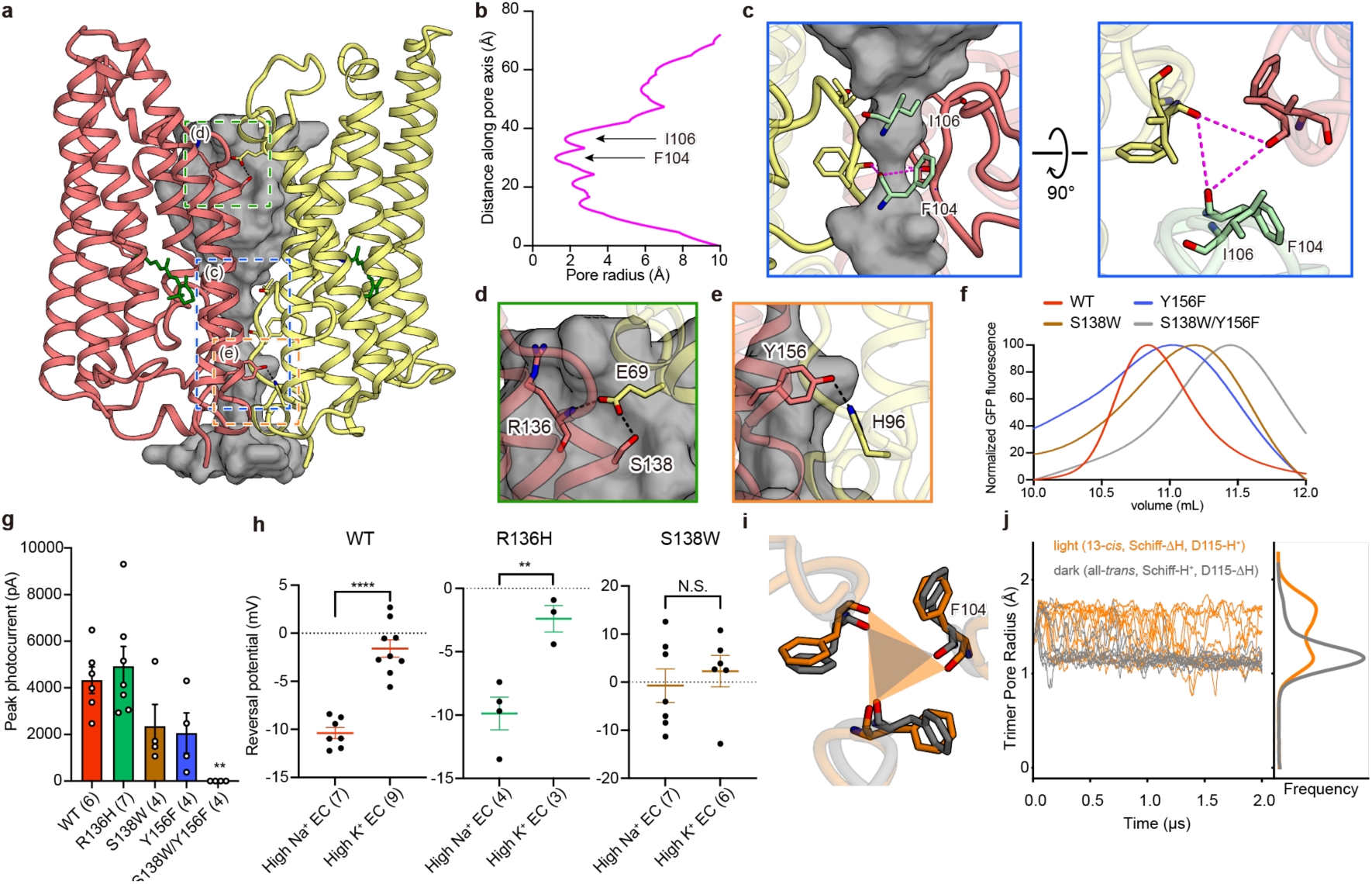
The hydrophilic pore within the trimer interface. **a**, Location of the pore within the trimer interface. The trimer pore pathway is depicted as grey mesh and only two protomers are shown for clarity. **b**, Pore radii as a function of the distance along the pore axis, calculated with HOLE. **c**, Magnified views of the blue-boxed region from **a**, the constriction formed by ECL1, from two angles. **d**, **e**, Magnified views of the green- and orange-boxed regions in **a**, the hydrogen bond interactions between protomers at the intracellular (**d**) and extracellular (**e**) side. **f**, Fluorescent size-exclusion chromatography (FSEC) traces of ChRmine WT, S138W, Y156F, and S138W/Y156F mutants, suggesting the destabilization of the trimer by mutations. **g**, Photocurrent amplitudes for the mutants shown in **f** as well as R136H (as a negative control). **h**, Reversal potentials of ChRmine WT, R136H, and S138W mutants, showing that S138W mutation modulates cation selectivity. Data are mean ± s.e.m; one-way ANOVA followed by Dunnett’s test for the middle and unpaired t-test for the right panel. \*\**P* < 0.01 and \*\*\*\**P* < 0.0001. Sample size (number of cells) indicated in parentheses. N.S., not significant. **i**, Visualization of the three F104 residues that define the trimer pore constriction site from representative frames of the dark and light state channels in simulation. **j**, Trimer pore radius was calculated for each of 10 independent 2 µs simulations. Pore radius is larger in light-state simulations than dark-state simulations (p < 0.001, Welch’s t-test).

To test this possibility, we first focused on the trimer interface. ChRmine exhibits three intermolecular hydrogen bond interactions between the adjacent protomers: S138 and E69, the main chain amide of R136 and E69, and Y156 and H96 (Fig. 4d and e). Therefore, we introduced mutations to each of these residues to destabilize trimer assembly. We used fluorescence-detection size exclusion chromatography (FSEC)^33^ to evaluate oligomerization, and found that S138W or Y156F mutation shifts the equilibrium to favor monomeric states, and S138W/Y156F double mutation almost completely dissociate the trimer into the monomeric state (Fig. 4f). Subsequent electrophysiology experiments revealed that these single and double mutations moderately and severely decrease channel activity, respectively (Fig. 4g and Extended Data Fig. 9), and S138W mutation additionally reduces overall ChRmine channel cation selectivity (Fig. 4h). Since S138 and Y156 are unequivocally located far from the canonical ion-conducting pathway within the monomer, and the R136H mutation (which does not affect protomer-protomer interactions) does not change channel activity or cation selectivity, these results support the novel idea that the trimer pore can serve as an auxiliary ion-conducting pathway.

To further test this hypothesis, we performed all-atom molecular dynamics simulations of ChRmine in either the dark state or the M intermediate (light state). In these simulations of the dark state, the retinal is left in the all-*trans* configuration and the Schiff base is protonated; in simulations of the light state, the retinal is isomerized to the 13-*cis* configuration and the Schiff base proton is transferred to the putative proton acceptor D115. Although simulations were not long enough to span complete activation of the channel, we did observe key early conformational changes along the pathway toward activation in the light state simulations, in which the trimer pore alternated between a wider, “open” state and a narrower, “closed” state; in contrast, for dark state simulations, the trimer pore remained in the closed state. The trimer pore radius (the radius of the constriction site formed by backbone interactions between the three F104 residues on each of the monomers) was significantly increased in light state simulations compared to dark state simulations (Fig. 4i and j), which sufficed to allow multiple water molecules to pass through the pore (Extended Data Fig. 11g and h). While the pore did not yet attain a radius sufficient for ion conduction over the timescale of our simulations, these results suggest that the trimer pore is cooperatively coupled to retinal isomerization and support the idea (consistent with the observed ion-selectivity change arising from mutation at the trimer pore (Fig. 4h) that the trimer pore can act as a novel secondary channel, via a structural mechanism not accessible to dimerizing chlorophyte channelrhodopsins or trimerizing pump rhodopsins (Supplementary discussion).

## Discussion

The unusual properties of the microbial channelrhodopsin ChRmine opened up new avenues of investigation for optogenetics in the study of cell-specific activity within biological systems^1, 3^. We also have found that ChRmine exhibits virtually no Ca^2+^ conductance (Extended Data Fig. 12), a property of substantial value in neuroscience applications for avoiding incidental induction of Ca^2+^ dependent plasticity.

Here, integrated structural, spectroscopic, electrophysiological, and computational analyses have revealed structure-function relationships underlying these remarkable properties, along with insights into the evolution of the microbial opsin gene families. First, despite its fundamentally distinct channel-based ion-conduction mechanism, we find that ChRmine is surprisingly similar to the *Hs*BR ion pump in terms of oligomerization number (Extended Data Fig. 7), overall monomeric structure (Fig. 1e), and proton acceptor (Fig. 2d), suggesting that ChRmine was evolved from archaeal ion-pumping rhodopsins. In *Hs*BR, D85 and D96 of DTD motif serve critical functions as relays for the pumping transfer of a single proton from the intracellular to extracellular side, in response to the absorbed photon. In ChRmine, these two Asps are conserved (D115 and D126) but not for proton-pumping relay purposes; rather, they form the two constriction sites (ICS and CCS) of the passive ion-conducting pore within the monomer. Thus, remarkably these two residues continue play critical roles, but in gating the ChRmine channel, rather than serving as relay points for proton transport in pumps (Supplementary discussion).

While ChRmine exhibits strong similarity to *Hs*BR, several unique features stand out in the high-resolution structure reported here (the first from a member of the BCCR family), including its long twisted ECL1. ECL1 not only significantly distorts the architecture around the Schiff base region but also increases the volume of the extracellular cavity within the monomer. In addition, the C-shaped part of ECL1 faces toward the center of the trimer and forms an ion selectivity filter-like structure along an unexpected central trimer pore. Interestingly, the length and sequence of ECL1 are not highly conserved among BCCRs, and cation selectivity has been reported to be different between BCCRs that we would predict to have long vs. short ECL1 domains^34^; this phylogenetic pattern may further suggest that the trimer pore is involved in cation conduction, supporting our direct observation of reversal-potential shift with trimer pore mutation (Fig. 4h).

To further deepen our understanding of the molecular mechanisms of light-gated ion conduction, additional studies (including structural determination of ChRmine in intermediate states) will be important (Supplementary discussion). However, our current structural information of ChRmine already provides a framework for further development of ChRmine-based optogenetic tools, and examples of such structure-guided engineering with application to neuroscience will be shown in accompanying work (manuscript in preparation). Additional insights into evolutionary and functional relationships among different members of the channelrhodopsin family will further arise from future structures of channelrhodopsin variants with different kinetic, spectral, and ion-selectivity properties.

During preparation of this manuscript, the deep learning-based protein-structure prediction program AlphaFold 2 was released^35^; we therefore generated the predicted structures (hereafter referred to as AF2 models) of ChRmine for comparison with our cryo-EM structure. Aspects of the overall structure of ChRmine could be predicted and the AF2 models could be superposed onto our cryo-EM structure (Extended Data Fig. 13a, b). However, striking differences were observed in N- and C-terminal regions, the cytoplasmic sides of TM5-7, and ICL1 and ECL1 (Extended Data Fig. 13c, d). In particular, the pLDDT (predicted local distance difference test) score^36, 37^ of most ECL1 was so low that no AF2 models could accurately predict the structure of ECL1 and the resulting pore within the trimer interface (Extended Data Fig. 13e). In addition, while D115 exhibited an exceptionally high pLDDT score (>90) in ECL1, its unique conformation was not reproduced by AF2 models; the models incorrectly predict the canonical conformation for D115, an outcome readily explicable due to biasing by the previously solved microbial rhodopsin structures deposited to the PDB database (Extended Data Fig. 8c and 13f). These results indicate that AlphaFold 2 is an outstanding program for structure prediction, but an experimentally-determined structure was clearly needed to discover and interpret the critical novel features of ChRmine, including the unique architecture of the Schiff base and an auxiliary ion-conducting pore within the trimer. A general cautionary note for the field is that when using the AF2 models, it would be important to carefully interpret the structures even if the pLDDT score is high.

Before this study, high-resolution structures of channelrhodopsins had been experimentally determined only using crystallography techniques. However, as shown in this study, we find that the combination of antibody and single-particle cryo-EM techniques is powerful enough to determine the high-resolution structure of small ion-transporting rhodopsins like ChRmine (in fact leading here to the highest-resolution cryo-EM structure of any rhodopsin yet reported, and the first in which some hydrogen atoms can be visualized), thus representing a new and promising option for structural analysis of microbial rhodopsins alongside X-ray crystallography. The two technologies, as well as structure prediction methods, may complement each other’s shortcomings and together accelerate the structural biology of microbial rhodopsins, and the resulting structural information will lead to both further development of optogenetics and basic mechanistic understanding of these remarkable photoreceptor proteins.

## Methods

### Cloning, Protein Expression, and Purification

Wild-type ChRmine was modified to include an N-terminal influenza hemagglutinin (HA) signal sequence and FLAG-tag epitope, and C-terminal enhanced green fluorescent protein (eGFP) and 10 × histidine tag; the N-terminal and C-terminal tags are removable by human rhinovirus 3C protease cleavage. The construct was expressed in *Spodoptera frugiperda* Sf9 insect cells using the pFastBac baculovirus system. Sf9 insect cells were grown in suspension to a density of 3.5 × 10^6^ cells/mL, infected with ChRmine baculovirus and shaken at 27°C for 24 h. Then, 10 µM all-*trans-*retinal (ATR) (Sigma) was supplemented to the culture and shaken continued for 24 more hours. The cell pellets were lysed with a hypotonic lysis buffer (20 mM HEPES pH 7.5, 20 mM NaCl, 10 mM MgCl_2_, 1 mM Benzamidine, 1 µg/ml leupeptin), and cell pellets were collected by centrifugation at 10,000 ×*g* for 30 min. The above process was repeated twice; then, cell pellets were disrupted by homogenizing with a glass dounce homogenizer in a hypertonic lysis buffer (20 mM HEPES pH 7.5, 1 M NaCl, 10 mM MgCl_2_, 1 mM Benzamidine, 1 µg/ml leupeptin), and crude membrane fraction was collected by ultracentrifugation. The above process was repeated twice; then, the membrane fraction was homogenized with a glass douncer in a solubilization buffer (1% *n*-dodecyl-β-D-maltoside (DDM), 0.2% cholesteryl hemisuccinate (CHS), 20 mM HEPES pH 7.5, 500 mM NaCl, 20% glycerol, 5 mM imidazole, 1 mM benzamidine, 1 µg/ml leupeptin) and solubilized for 2 h in 4°C. The insoluble cell debris was removed by centrifugation (125,000 ×*g*, 1 h), and the supernatant was mixed with the Ni-NTA superflow agarose resin (QIAGEN) for 1 h in 4°C. The Ni-NTA resin was collected into a glass chromatography column, washed with wash 1 buffer (0.05% DDM, 0.01% CHS, 20 mM HEPES pH7.5, 100 mM NaCl, 50 mM imidazole), wash 2 buffer (0.05% DDM, 0.06% GDN (glyco-diosgenin), 0.016% CHS, 20 mM HEPES pH7.5, 100 mM NaCl, 50 mM imidazole), and wash 3 buffer (0.06% GDN, 0.006% CHS, 20 mM HEPES pH7.5, 100 mM NaCl, 50 mM imidazole), and was eluted in a wash 3 buffer supplemented with 300 mM imidazole. After cleavage of the FLAG tag and eGFP-His_10_ tag His-tagged 3C protease, the sample was reloaded onto the Ni-NTA column to capture the cleaved eGFP-His_10_. The flow-through containing ChRmine was collected, concentrated, and purified through gel-filtration chromatography.

### Antibody generation

All the animal experiments conformed to the guidelines of the Guide for the Care and Use of Laboratory Animals of Japan and were approved by the Kyoto University Animal Experimentation Committee. Mouse monoclonal antibodies against ChRmine were raised according to previously-described methods^38^. Briefly, a proteoliposome antigen was prepared by reconstituting purified, functional ChRmine at high density into phospholipid vesicles consisting of a 10:1 mixture of chicken egg yolk phosphatidylcholine (egg PC; Avanti Polar Lipids) and the adjuvant lipid A (Sigma-Aldrich) to facilitate immune response. BALB/c mice were immunized with the proteoliposome antigen using three injections at two-week intervals. Antibody-producing hybridoma cell lines were generated using a conventional fusion protocol. Biotinylated proteoliposomes were prepared by reconstituting ChRmine with a mixture of egg PC and 1,2-dipal-mitoyl-sn-glycero-3-phosphoethanolamine-N-(cap biotinyl) (16:0 biotinyl Cap-PE; Avanti), and used as binding targets for conformation-specific antibody selection. The targets were immobilized onto streptavidin-coated microplates (Nunc). Hybridoma clones producing antibodies recognizing conformational epitopes in ChRmine were selected by an enzyme-linked immunosorbent assay on immobilized biotinylated proteoliposomes (liposome ELISA), allowing positive selection of the antibodies that recognized the native conformation of ChRmine. Additional screening for reduced antibody binding to SDS-denatured ChRmine was used for negative selection against linear epitope-recognizing antibodies. Stable complex formation between ChRmine and each antibody clone was checked using fluorescence-detection size-exclusion chromatography. The sequence of the Fab from the antibody clone number YN7002_7 (named Fab02) was determined via standard 5’-RACE using total RNA isolated from hybridoma cells.

### Formation and purification of the ChRmine-Fab02 complex

Purified ChRmine was mixed with a fourfold molar excess of Fab, and the coupling reaction proceeded at 4°C overnight. The ChRmine-Fab02 complex was purified by size exclusion chromatography on a Superdex 200 increase 10/300 GL column (Cytiva).

### Cryo-EM data acquisition and image processing

Cryo-EM images were acquired at 300 kV on a Krios G3i microscope (Thermo Fisher Scientific) equipped with a Gatan BioQuantum energy filter and a K3 direct detection camera in the electron counting mode. The movie dataset was collected in a correlated double sampling (CDS) mode, using a nine-hole image shift strategy in the SerialEM software^38^, with a nominal defocus range of 0.8 to 1.6 μm. The 3,528 movies were acquired at a dose rate of 6.3 e^-^/pixel/s, at a pixel size of 0.83 Å and a total dose of 46 e^-^/Å^2^.

Image processing was performed in RELION-3.1^39^. Beam-induced motion correction and dose weighting were performed with RELION’s implementation of the MotionCor2 algorithm^40^, and CTF parameters were estimated with CTFFIND-4.1.13^40^. Particles were first picked using the Laplacian-of-gaussian algorithm, and 2D class average images were generated as templates for reference-based auto-picking. Reference-based picked 2,958,159 particles were subjected to several rounds of 2D and 3D classifications. The selected 555,801 particles were subjected to a 3D auto-refinement, resulting in a 2.8 Å map. Subsequently, Bayesian polishing^41^ and CTF refinement^42^, followed by a 3D auto-refinement, resulted in a 2.6 Å map. Micelle and constant regions of Fab fragments densities were subtracted from particle images, and the subtracted particles were subjected to a masked 3D classification without alignment. After a 3D auto-refinement of selected 185,895 particles, three runs of CTF refinement were performed in order as follows: refining magnification anisotropy; refining optical aberrations; refining per-particle defocus and per-micrograph astigmatism. Another round of 3D auto-refinement yielded a 2.13 Å map. These particles were subjected to, a second round of Bayesian polishing, CTF refinement and a transmembrane region-focused 3D auto-refinement with the reconstruction algorithm SIDESPLITTER^43^, resulting in the final map at a global resolution of 2.03 Å.

### Model building and refinement

An initial model was formed by rigid body fitting of the C1C2 (PDB ID: 3UG9)^18^. This starting model was then subjected to iterative rounds of manual and automated refinement in Coot^44^ and Refmac5^45^ in Servalcat pipeline^22^, respectively. The Refmac5 refinement was performed with the constraint of C3 symmetry. The final model was visually inspected for general fit to the map, and geometry was further evaluated using Molprobity^46^. The final refinement statistics for both models are summarized in Supplementary Table 2. All molecular graphics figures were prepared with UCSF Chimera^47^, UCSF ChimeraX^48^, and Cuemol (http://www.cuemol.org).

### High performance liquid chromatography (HPLC) analysis of retinal isomers

The retinal isomers were analyzed with an HPLC system equipped with a silica column (particle size 3 μm, 150 × 6.0 mm; Pack SIL, YMC, Japan), a pump (PU-4580, JASCO, Japan) and a UV–Visible detector (UV-4570, JASCO, Japan). The purified sample in a buffer containing 20 mM HEPES-NaOH (pH 7.5), 100 mM NaCl, 0.035% GDN, 0.0035% CHS (GDN:CHS = 10:1) were dark-adapted for two days at 4 °C. A 75 μL sample and 280 μL of 90% (v/v) methanol aqueous solution were mixed on ice and then 25 μL of 2 M hydroxylamine (NH_2_OH) was added to convert retinal chromophore into retinal oxime, which was extracted with 800 μL of *n*-hexane. A 200 μL of the extract was injected into the HPLC system. The solvent containing 15% ethyl acetate and 0.15% ethanol in hexane was used as a mobile phase at a flow rate of 1.0 mL min^-1^. Illumination was performed on ice with green light (530 ± 5 nm) for 20 s for samples under illumination and 60 s for light adaptation. The molar composition of the sample was calculated from the areas of the peaks and the molar extinction coefficients at 360 nm (all-*trans*-15-*syn*: 54,900 M^-1^cm^-1^; all-*trans*-15-*anti*: 51,600 M^-1^cm^-1^; 13-*cis*-15-*syn*, 49,000 M^-1^cm^-1^; 13-*cis*-15-*anti*: 52,100 M^-1^cm^-1^; 11-*cis*-15-*syn*: 35,000 M^-1^cm^-1^; 11-*cis*-15-*anti*: 29,600 M^-1^cm^-1^)^49^.

### Preparation of lipid-reconstituted ChRmine

For HS-AFM imaging of lipid-reconstituted ChRmine, we applied membrane scaffolding proteins (MSP), which were developed for nanodisc technology^50, 51^. We followed the manufacturer’s protocol for the nanodisc (Sigma-Aldrich, St. Louis, MO, USA) with minor modifications as described previously^52^. Briefly, for reconstituted lipids, we used a mixture of phospholipids, asolectin from soybean (Sigma-Aldrich, No. 11145). Asolectin (120 μg) was dissolved in chloroform and then evaporated under N_2_ gas to completely remove the solvent. Then, the lipids were suspended in 50 μL buffer A (20 mM HEPES-KOH (pH 7.4), 100 mM NaCl, and 4% DDM) and sonicated for ∼1 min with a tip-sonicator. Next, dissolved membrane proteins (1 nmol) and MSP (50 μL, 1 mg/mL) (MSP1E3D1, Sigma-Aldrich, No. M7074) were added to the lipid suspension and mixed for ∼1 h while rotating in the dark at 4°C. Finally, we added 60 mg Bio-beads SM-2 (Bio-Rad, Hercules, CA, USA, No. 1523920) and dialyzed the samples in detergent overnight at 4°C. According to the manufacturer’s protocol, nanodisc samples should be fractionated on a column to purify the nanodiscs based on size (∼10 nm in diameter). Here, we did not purify the reconstituted samples, but obtained flat membranes with limited sizes <30 nm in diameter.

### High-speed AFM measurements

A homemade HS-AFM operated in tapping mode was used^52, 53^. An optical beam deflection detector detected the cantilever (Olympus, Tokyo, Japan: BL-AC10DS-A2) deflection using an infrared (IR) laser at 780 nm and 0.7 mW. The IR beam was focused onto the back side of the cantilever covered with a gold film through a ×60 objective lens (Nikon, Tokyo, Japan: CFI S Plan Fluor ELWD 60x). The reflected IR beam was detected by a two-segmented PIN photodiode. The free oscillation amplitude of the cantilever was ∼1 nm and set-point amplitude was approximately 90% of the free amplitude for feedback control of HS-AFM observation. An amorphous carbon tip (∼500 nm length), grown by electron beam deposition by scanning electron microscope, was used as an AFM probe. As a HS-AFM substrate, a mica surface treated with 0.01% (3-aminopropyl) triethoxysilane (Shin-Etsu Silicone, Tokyo, Japan) was used. All HS-AFM experiments were carried out in buffer solution containing 20 mM Tris–HCl (pH 8.0) and 100 mM NaCl at room temperature (24–26 °C) and data analyses were conducted using laboratory-developed software based on IgorPro 8 software (WaveMetrics, USA). We usually used a scan area of 43 × 32 nm^2^ with 130 × 95 pixels. HS-AFM images were captured at frame rates of 2 fps. All HS-AFM images were processed by Gaussian noise-reduction filters.

### Measurement of UV absorption spectra

Protein absorbance spectra were measured with an Infinite M1000 microplate reader (Tecan Systems Inc.) using 96 well plates (Thermofisher scientific). The ChRmine samples were suspended in a buffer containing 100 mM NaCl, 0.05% DDM, 0.01% CHS, and 20 mM sodium citrate, sodium acetate, sodium cacodylate, HEPES, Tris, CAPSO, or CAPS. pH was adjusted from 4 to 10 by addition of NaOH or HCl.

### Laser flash photolysis

For the laser flash photolysis spectroscopy, wildtype ChRmine was solubilized in 20 mM HEPES-NaOH (pH 7.5), 100 mM NaCl, 0.035% GDN, 0.0035% CHS (GDN:CHS = 10:1) or 20 mM Acetate (pH 4.0), 100 mM NaCl, 0.03% GDN, 0.003% CHS (GDN:CHS = 10:1), and ChRmine D115N and D253N mutants were solubilized in 20 mM Acetate (pH 4.0), 100 mM NaCl, 0.03% GDN, 0.003% CHS (GDN:CHS = 10:1). The optical density of the protein solution was adjusted to ∼0.4 (protein concentration ∼0.28 mg/mL) at the absorption maximum wavelengths. The laser flash photolysis measurements were conducted as previously described^17^. ChRmine wildtype at pH 7.5 was excited by the second harmonics of a nanosecond-pulsed Nd^3+^-YAG laser (excitation wavelength (*λ*_exc_) = 532 nm, 4.5 mJ/pulse, 1.4–0.5 Hz, INDI40, Spectra-Physics, CA), and nano-second pulse from an optical parametric oscillator (4.5 mJ/pulse, basiScan, Spectra-Physics, CA) pumped by the third harmonics of Nd^3+^-YAG laser (*λ* = 355 nm, INDI40, Spectra-Physics, CA) was used for the excitation of ChRmine wildtype at pH 4.0 (*λ*_exc_ = 505 nm), ChRmine D115N (*λ*_exc_ = 488 nm) and D253N (*λ*_exc_ = 500 nm). Transient absorption spectra were obtained by monitoring the intensity change of white-light from a Xe-arc lamp (L9289-01, Hamamatsu Photonics, Japan) passed through the sample with an ICCD linear array detector (C8808-01, Hamamatsu, Japan). To increase the signal-to-noise (S/N) ratio, 45–60 spectra were averaged, and the singular-value-decomposition (SVD) analysis was applied. To measure the time-evolution of transient absorption change at specific wavelengths, the light of Xe-arc lamp (L9289-01, Hamamatsu Photonics, Japan) was monochromated by monochromators (S-10, SOMA OPTICS, Japan) and the change in the intensity after the photo-excitation was monitored with a photomultiplier tube (R10699, Hamamatsu Photonics, Japan). To increase S/N ratio, 100–200 signals were averaged. The time-evolution of transient absorption change was analyzed by global multi-exponential fitting to determine the time constant of each reaction step and absorption spectra of the photo-intermediates. Some reaction steps were reproduced by double or triple exponentials. In this case, the averaged time constant calculated by

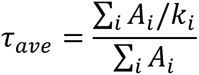

where *A*_i_ and *k*_i_ are the amplitude at a wavelength of representing the M-intermediate and the rate constant of i-th exponential function (i = 1–3).

### MD simulation system setup

We performed simulations of ChRmine trimer in different modeled photoactivation states. In the dark state, the Schiff base remained protonated and in an all-*trans* configuration; residue D115 was unprotonated. In the light state the Schiff base was deprotonated and in a 13-*cis*, 15-*anti* retinal configuration; residue D115 was protonated. We initiated simulations from the structures presented in this paper. Neutral acetyl and methylamide groups were added to cap the N- and C-termini, respectively of protein chains. Titratable residues were kept in their dominant protonation states at pH 7, except for D115 which was protonated for light state simulations to simulate proton transfer. Histidine residues were modeled as neutral, with a hydrogen bound to either the delta or epsilon nitrogen depending on which tautomeric state optimized the local hydrogen-bonding network. We performed 10 independent simulations in which initial atom velocities were assigned randomly and independently for four separate conditions: with all three monomers in the dark state, with one monomer activated in the light state and two in the dark state, with two monomers activated in the light state and one in the dark state and finally with all three monomers in the light state.

Dowser was used to add water molecules to protein cavities and aligned on the entire trimer structure. The aligned structures were inserted into a pre-equilibrated palmitoyl-oleoyl-phosphatidylcholine (POPC) membrane bilayer using Dabble^54^. Sodium and chloride ions were added to neutralize each system at a concentration of 150 mM. Systems were composed of ∼100,200 atoms, including ∼215 lipid molecules and ∼19,000 water molecules. Approximate system dimensions were 80 Å x 90 Å x 85 Å.

### MD simulation protocol

The systems were first heated over 12.5 ps from 0 K to 100 K in the NVT ensemble using a Langevin thermostat with harmonic restraints of 10.0 kcal·mol^-1^ ·Å-2 on the non-hydrogen atoms of the lipids, protein, and ligand. Initial velocities were sampled from a Boltzmann distribution. The systems were then heated to 310 K over 125 ps in the NPT ensemble. Equilibration was performed at 310 K and 1 bar in the NPT ensemble, with harmonic restraints on the protein and ligand non-hydrogen atoms tapered off by 1.0 kcal·mol^-1^ ·Å^-2^ starting at 5.0 kcal·mol^-1^ ·Å^-2^ in a stepwise manner every 2 ns for 10 ns, and finally by 0.1 kcal·mol^-1^ ·Å^-2^ every 2 ns for an additional 18 ns. To generate light state simulations an additional torsional restraint step of 2.7 ps was added to flip the retinal to the 13-*cis*, 15-*anti* configuration.

All restraints were removed during production simulation. Production simulations were performed at 310 K and 1 bar in the NPT ensemble using the Langevin thermostat and Monte Carlo barostat. The simulations were performed using a timestep of 4.0 fs while employing hydrogen mass repartitioning^55^. Bond lengths were constrained using SHAKE^56^. Non-bonded interactions were cut off at 9.0 Å, and long-range electrostatic interactions were calculated using the particle-mesh Ewald (PME) method with an Ewald coefficient (β) of approximately 0.31 Å and B-spline interpolation of order 4. The PME grid size was chosen such that the width of a grid cell was approximately 1 Å. We employed the CHARMM36m force field for protein molecules, the CHARMM36 parameter set for lipid molecules and salt ions, and the associated CHARMM TIP3P model for water^57–59^. All simulations were performed on the Sherlock computing cluster at Stanford University.

### MD simulation analysis protocol

The AmberTools17 CPPTRAJ package^60^ was used to reimage trajectories, while Visual Molecular Dynamics (VMD)^61^ was used for visualization and analysis.

Computation of the trimer pore radius and monomer pore radius was conducted with HOLE, a Monte Carlo method that identifies the pore through a protein channel^62^. The minimum pore radius was identified for each frame of each simulation. Pore analysis was completed using the entire trimer for the trimer pore radius, and for the monomer pore radius the pore was identified from a subselection of each monomer (residue numbers 253 to 259, 113 to 120, 78 to 83, and 38 to 43) to target the section of the protein near the retinal and proton acceptor D115.

### In vitro Electrophysiology

HEK293 cells (authenticated by the vendor, not tested for mycoplasma contamination) expressing ChRmine variants were recorded as previously described^30, 63^. Briefly, after 24–48 h of transfection, cells were placed in an extracellular tyrode medium (150 mM NaCl, 4 mM KCl, 2 mM CaCl_2_, 2 mM MgCl_2_, 10 mM HEPES pH 7.4, and 10 mM glucose). Borosilicate patch pipettes (Harvard Apparatus) with resistance of 4 – 6 MOhm were filled with intracellular medium (140 mM potassium-gluconate, 10 mM EGTA, 2 mM MgCl_2_ and 10 mM HEPES pH 7.2). Light was delivered with the Spectra X Light engine (Lumencor) connected to the fluorescence port of a Leica DM LFSA microscope with a 580 nm filter for orange light generation.

Channel kinetics and photocurrent amplitudes were measured in voltage clamp mode at -70 mV holding potential. To determine channel kinetics and photocurrent amplitudes, traces were first smoothed using a lowpass Gaussian filter with a −3 dB cutoff for signal attenuation and noise reduction at 1,000 Hz and then analysed in Clampfit software (Axon Instruments). Liquid junction potentials were corrected using the Clampex built-in liquid junction potential calculator as previously described. Statistical analysis was performed with t-test or one-way ANOVA, and the Kruskal–Wallis test for non-parametric data, using Prism 7 (GraphPad) software. Data collection across opsins was randomized and distributed to minimize across-group differences in expression time, room temperature, and related experimental factors.

### Ion Selectivity Testing

Cells and devices for the measurement were prepared as described in the previous section. For the high sodium extracellular / high potassium intracellular condition, we used sodium bath solution containing 120 mM NaCl, 4 mM KCl, 2 mM CaCl_2_, 2 mM MgCl_2_, and 10 mM HEPES pH 7.2 (with glucose added up to osm 310 mOsm), along with potassium pipette solution containing 120 mM KCl, 10 mM EGTA, 4 mM NaCl, 2 mM CaCl_2_, 2 mM MgCl_2_, and 10mM HEPES pH 7.2 (with glucose added up to osm ∼290). For the high potassium extracellular / high sodium intracellular condition, NaCl and KCl concentrations were reversed, and all other ionic concentrations were kept constant. For ion selectivity measurements, ions in both bath and pipette solutions were replaced with either 120 mM NaCl, 120 mM KCl, 80 mM CaCl_2_, 80 mM MgCl_2_ or 120 mM NMDG-Cl, with all other components at low concentrations (4 mM NaCl, 4 mM KCl, 2 mM CaCl_2_, 2 mM MgCl_2_, and 10mM HEPES). Glucose was added to increase intracellular solution to 310 mOsm and extracellular solution to 290 osm. Photocurrent amplitudes were measured at -70 mV holding membrane potential. Equilibrium potentials were measured by holding membrane potentials from -75 mV to + 45 mV in steps of 10 mV.

### Data availability

The raw images of ChRmine after motion correction has been deposited in the Electron Microscopy Public Image Archive, under accession EMPIAR-xxxxx. The cryo-EM density map and atomic coordinates for ChRmine have been deposited in the Electron Microscopy Data Bank and PDB, under accessions EMD-xxxxx and xxxx, respectively. The cryo-EM density map for ChRmine-Fab02 complex has been deposited in the Electron Microscopy Data Bank under accessions EMD-xxxxx. All other data are available upon request to the corresponding authors.

## Acknowledgements

We thank Radostin Danev and Masahide Kikkawa (University of Tokyo) for setting up the cryo-EM infrastructure; Abhay Kotecha (Thermo Fisher Scientific) for the support of cryo-EM data collection; Andrés Maturana (Nagoya University) for the support of electrophysiology; and Kumika Hasegawa (University of Tokyo) and Cynthia Delacruz (Stanford University) for technical and administrative support of the project. This work was supported by the Medical Research Council, as part of UK Research and Innovation (MC_UP_A025_1012 to K.Y.), Osamu Hayaishi Memorial Scholarship for Study Abroad (M.I.), The Nakajima Foundation (H.E.K.), the Yamada Science Foundation (H.E.K.), the UTEC-UTokyo FSI Research Grant Program (H.E.K.), the Platform Project for Supporting Drug Discovery and Life Science Research (Basis for Supporting Innovative Drug Discovery and Life Science Research (BINDS)) from AMED under Grant Number JP18am0101079 (support no. 2165) and JP20am0101115 (support no. 2841), JSPS KAKENHI (JP20K21383 and JP21H01875 to K.I.), Japan Science and Technology Agency (JST) PRESTO (JPMJPR1782 to H.E.K. and JPMJPR1888 to T.N.), JST FOREST (JPMJFR204S to H.E.K.), the National Science Foundation NeuroNex program (K.D.), the NOMIS Foundation (K.D.), the Else Kröner Fresenius Foundation (K.D.), the Gatsby Foundation (K.D.), and a grant for channelrhodopsin structure determination from the NIMH (R01MH075957 to K.D.).

## Contributions

K.E.K. and M.F. performed the molecular cloning, expressed and purified the proteins, and prepared cryo-EM grids. T.K. obtained cryo-EM images, using the cryo-EM facility platform developed by O.N.. T.K. and T.E.M. processed cryo-EM data to generate 3D maps. K.E.K., K.Y., and H.E.K. built the model and refined the structure. Y.S.K. and Y.W. performed the electrophysiological analyses using HEK293 cells. E.T. and J.M.P. performed and analyzed the molecular dynamics simulations under the supervision of R.O.D. E.F.X.B. and Y.S.K. screened constructs and performed crystallography at an early stage. E.F.X.B. generated the predicted structures using AlphaFold 2. C.R. generated the cell cultures and carried out the molecular cloning and mutagenesis for electrophysiology. T.B. and M.I. provided input on electrophysiological considerations. K.E.K. measured UV-Vis absorption spectra. M.K. and K.I. performed flash photolysis experiments. T.N. performed HPLC analysis to determine the retinal isomer. M.S. performed HS-AFM experiments. T.U. and K.L. produced the Fab fragment that recognizes ChRmine, using the antibody production platform for membrane proteins developed by S.I. and N.N.. K.E.K., Y.S.K., K.D., and H.E.K. wrote the manuscript with input from all the authors. K.D. and H.E.K. supervised all aspects of the research.

---

**Extended Data Fig. 1.**
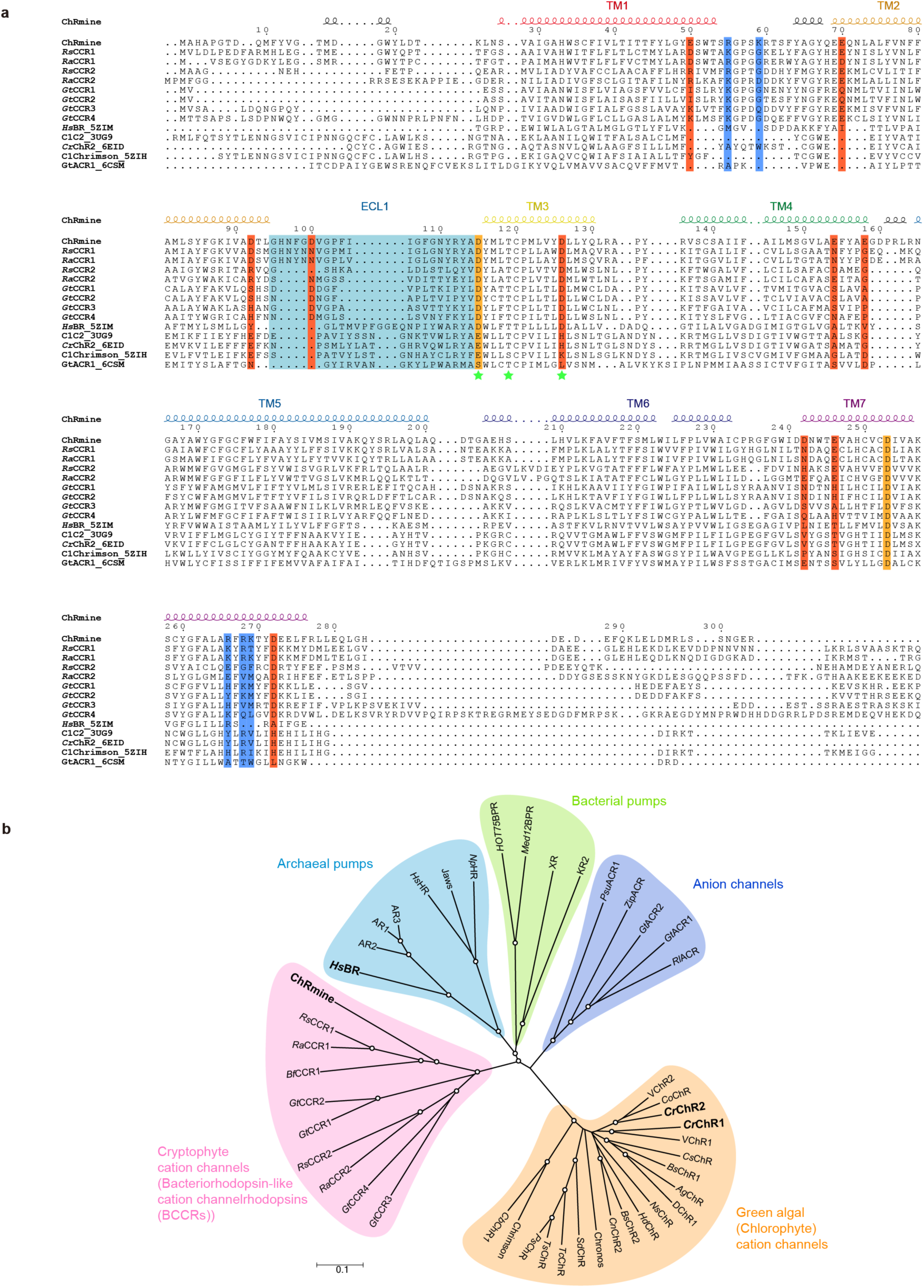
Structure-based sequence alignment and phylogenetic tree. The sequences are ChRmine (GenBank accession QDS02893.1), *Ra*CCR1 (GenBank QIU80793.1), *Rs*CCR1 (GenBank QIU80800.1), *Bf*CCR1 (GenBank QIU80783.1), *Gt*CCR1 (GenBank ANC73520.1), *Gt*CCR2 (GenBank ANC73518.1), *Gt*CCR3 (GenBank ANC73519.1), *Gt*CCR4 (GenBank ARQ20888.1), *Hs*BR (PDB ID: 5ZIM)^64^, C1C2 (PDB ID: 3UG9)^18^, *Cr*ChR2 (PDB ID: 6EID)^65^, *C1*Chrimson (PDB ID: 5ZIH)^66^, *Gt*ACR1 (PDB ID: 6CSM)^63^, *Cr*ChR1 (GenBank AAL08946.1), VChR1 (GenBank ABZ90900.1), VChR2 (GenBank ABZ90902.1), Chronos (GenBank KF992040.1), *Gt*ACR2 (GenBank AKN63095.1), *Rl*ACR (GenBank APZ76712.1), *Hs*HR (PDB ID: 1E12)^24^, BPRMed12 (PDB ID: 4JQ6)^67^, XR (PDB ID: 3DDL)^68^, KR2 (PDB ID: 3X3B)^69^. **a**, The sequence alignment was created using PLOMALS3D^70^ and ESPript 3^71^ servers. Secondary structure elements for ChRmine are shown as coils. Positive and negative charged residues are highlighted in blue and red, respectively. Green stars represent the DTD motif. The ECL1 of ChRmine is coloured light blue. The counterions are coloured orange. **b**, An unrooted phylogenetic tree was drawn for representative microbial rhodopsins using the Neighbor-Joining method^72^, and 1,000 bootstrap replicates. Evolutionary analyses were conducted in MEGA7^73^. White circles represent bootstrap values >85%.

**Extended Data Fig. 2.**
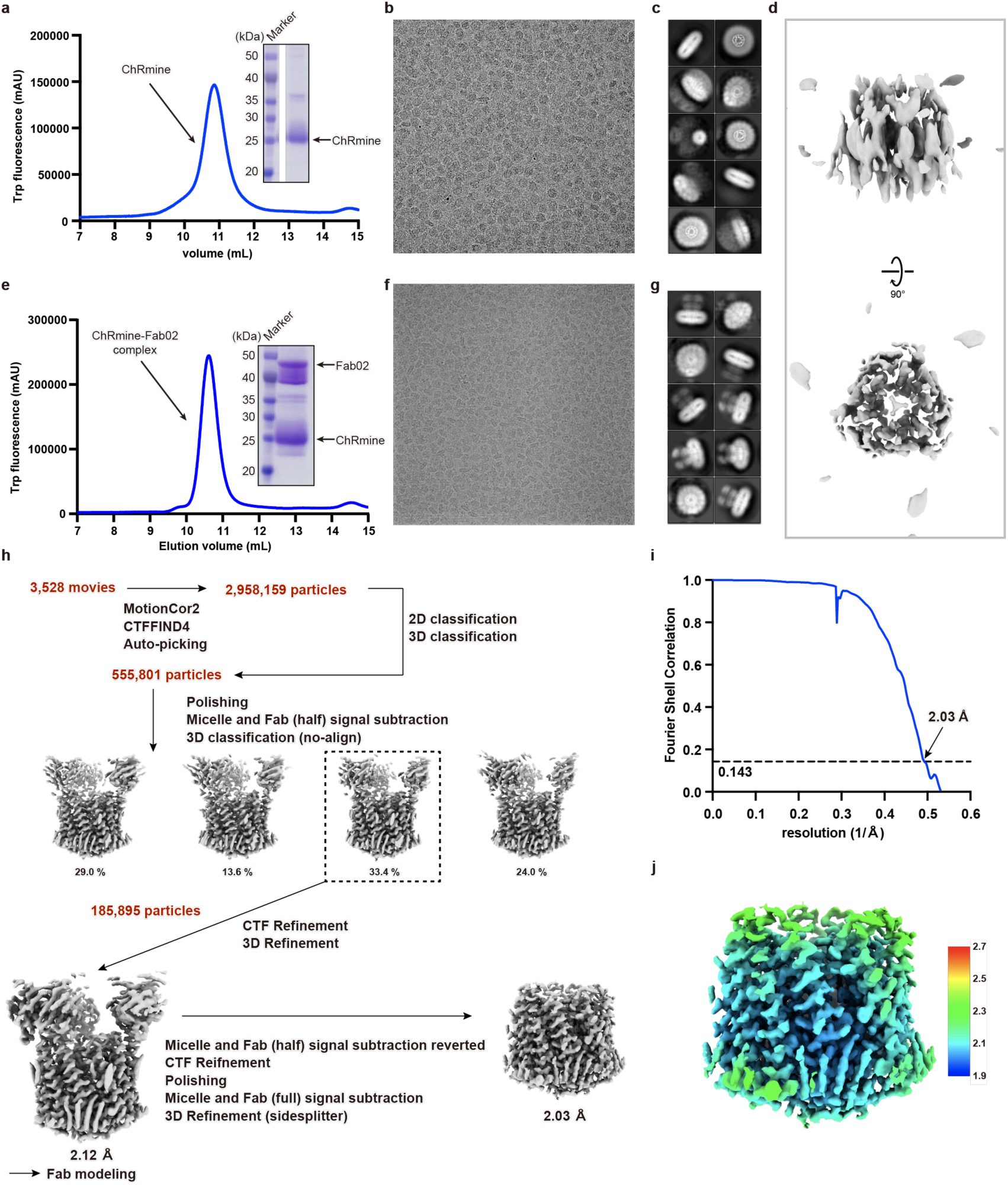
Biochemistry and cryo-EM of ChRmine and ChRmine-Fab02 complex. **a-d,** Panels corresponding to ChRmine alone. **e-j,** Panels corresponding to the ChRmine-Fab02 complex. **a,e**, Representative SEC trace with SDS-PAGE as inset. **b,f**, Representative cryo-EM micrograph. **c,g**, 2D-class averages. **d**, Low-resolution reconstruction of ChRmine. **h**, Data processing workflow of ChRmine-Fab02 complex. **i**, ‘Gold-standard’ FSC curves. **j**, Final cryo-EM map coloured by local resolution.

**Extended Data Fig. 3.**
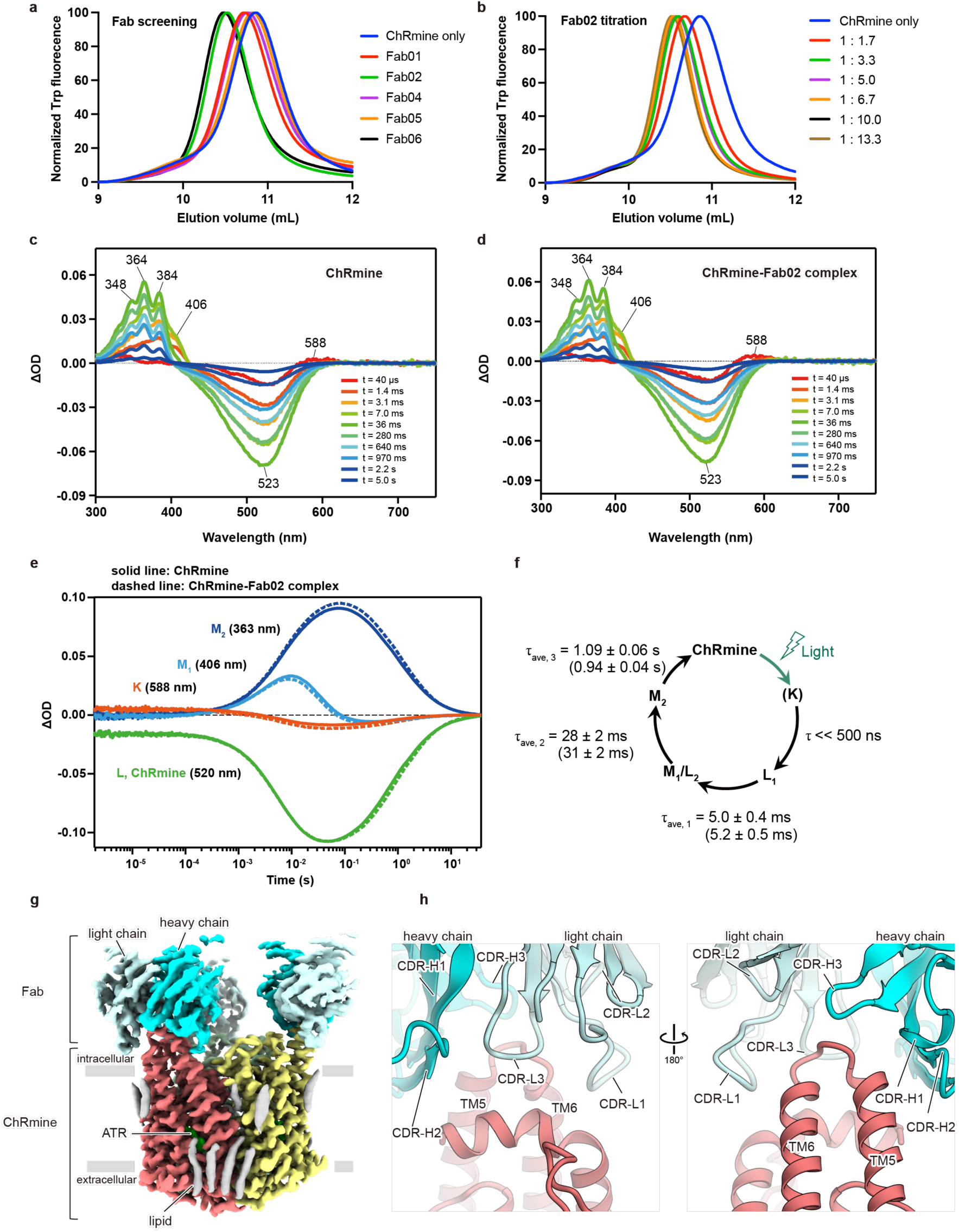
Spectroscopic and structural analysis of the ChRmine-Fab02 complex. **a**, FSEC screening of Fab fragments. **b**, Titration of Fab02 fragment against ChRmine. **c**,**d**, Transient absorption spectra of ChRmine WT (**c**) and the ChRmine-Fab02 complex (**d**) excited at *λ*_exc_ = 532 nm. **e**, Time series traces of absorption changes of ChRmine WT (solid line) and the ChRmine-Fab02 complex (right) at 363 (blue), 406 (cyan), 520 (green), 588 nm (red) probe wavelengths. **f**, Photocycle scheme of ChRmine determined by flash photolysis shown in **e**. **g**, Cryo-EM map of ChRmine-Fab02 complex. **h**, Interactions between ChRmine and Fab02.

**Extended Data Fig. 4.**
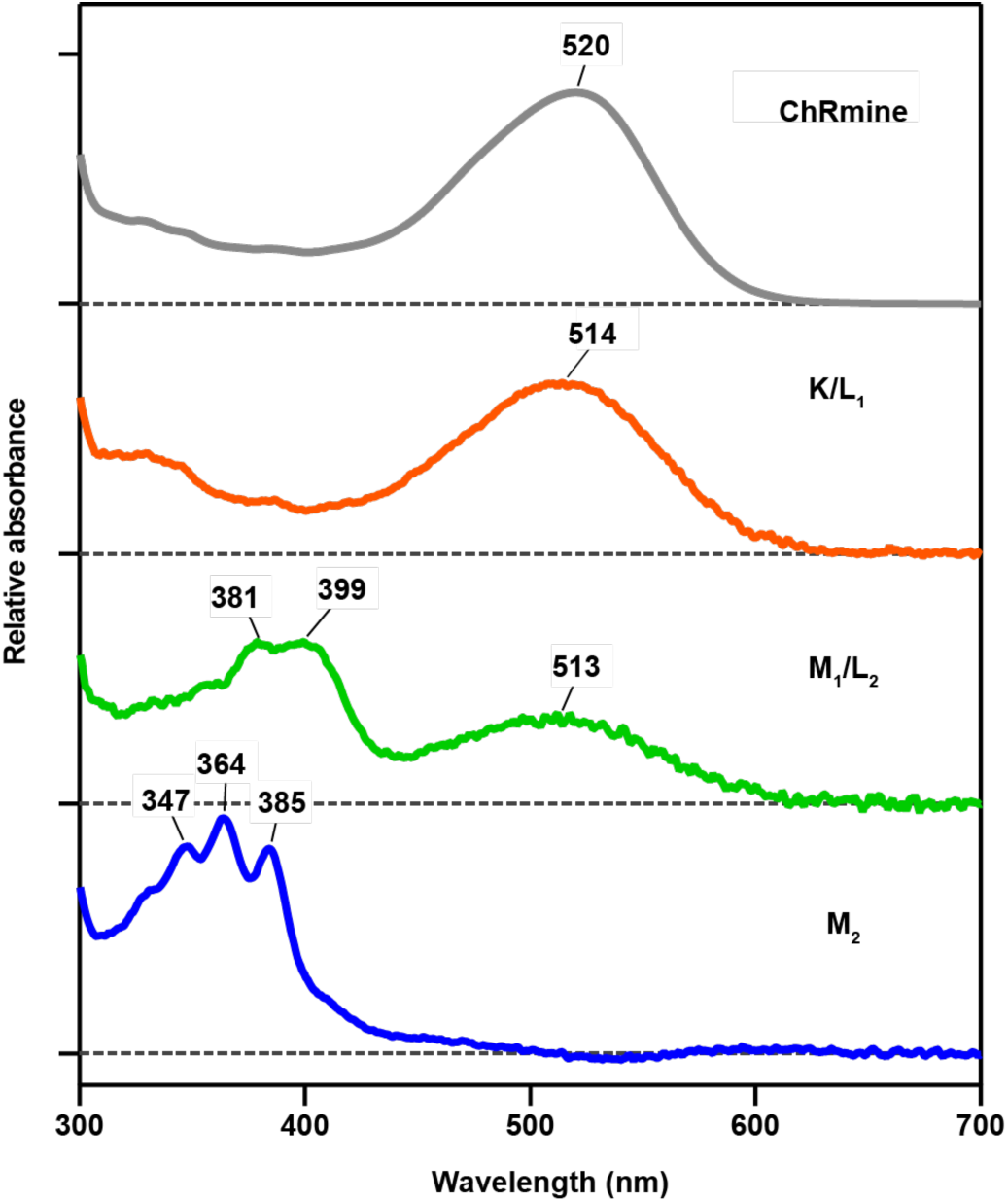
Relative absorption spectra of the initial state and each intermediate of ChRmine. The absorption spectra of the initial state (grey), L_1_ (orange), M_1_/L_2_ (green), and M_2_ (blue) of ChRmine calculated from the decay-associated-spectra of transient absorption changes^17^.

**Extended Data Fig. 5.**
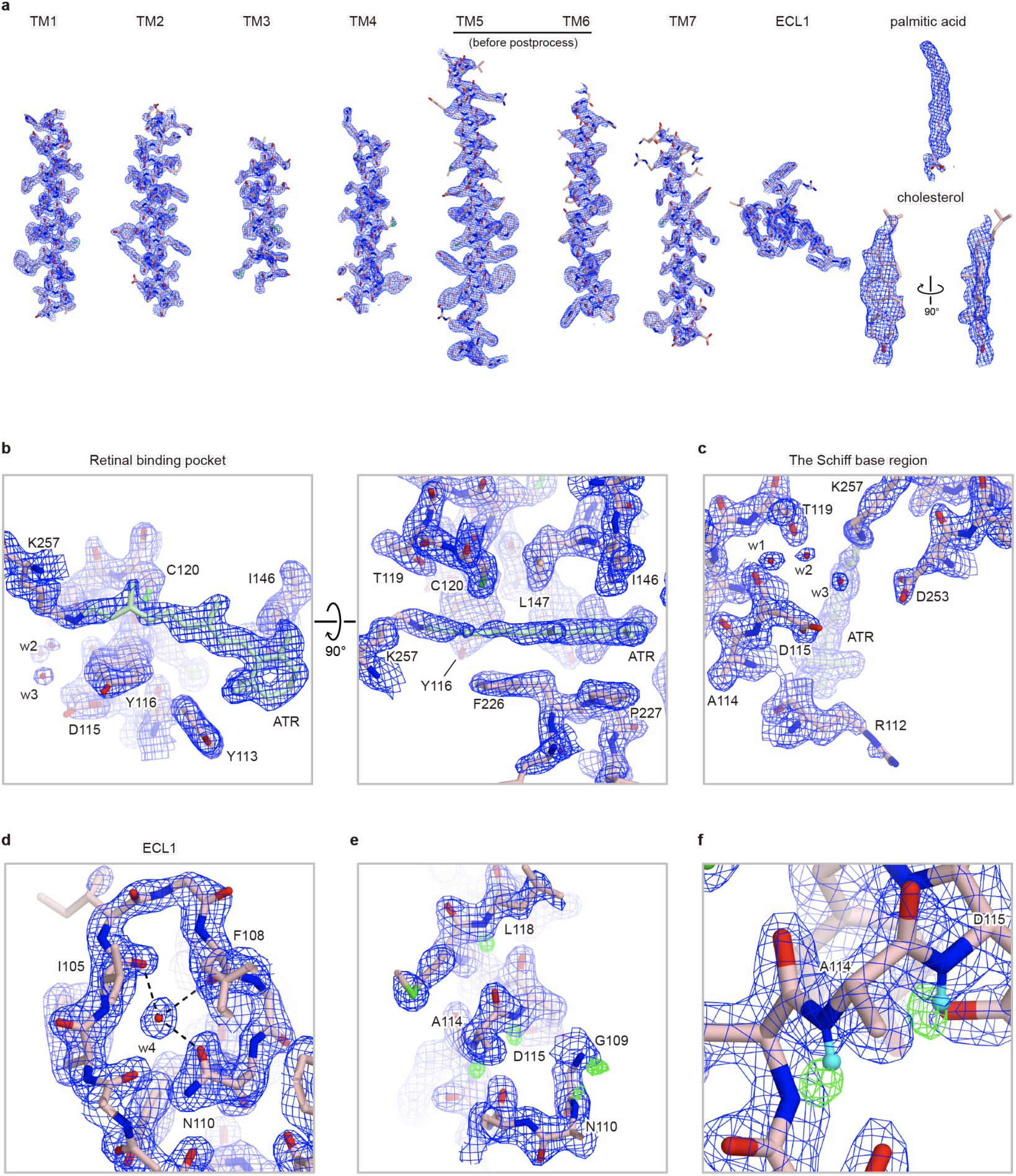
Cryo-EM density. **a-d**, Density (FSC-weighted sharpened map calculated by RELION3.1.1) and model for ChRmine, lipids (**a**), the retinal binding pocket (**b**), the Schiff base region (**c**), twisted ECL1 (**d**) **e,f**, Density and model near ECL1 region. FSC-weighted sharpened map calculated by RELION3.1.1 (blue) and *F_o_-F_c_* map calculated by Servalcat (green). Positive *F_o_-F_c_* difference densities (4.3σ, where σ is the standard deviation within the mask) are observed near nitrogen atoms, suggesting that these densities represent hydrogen atoms.

**Extended Data Fig. 6.**
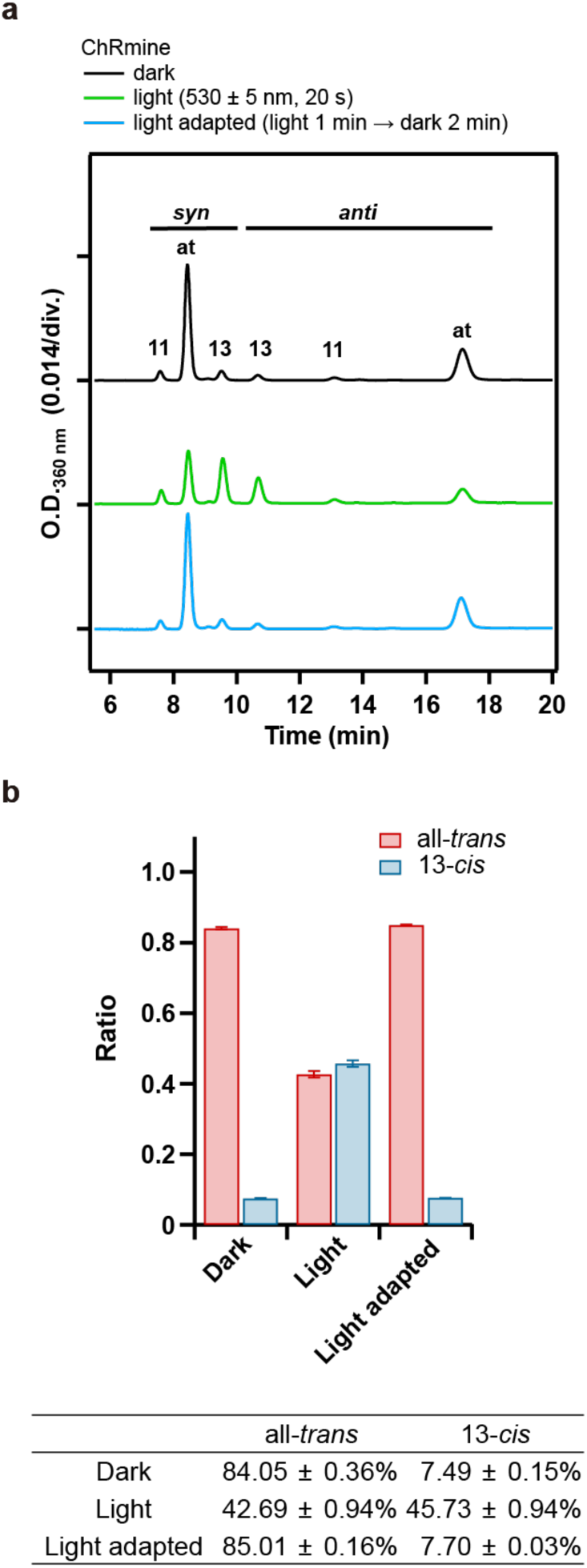
HPLC analysis of the chromophore configuration of ChRmine. **a**, Representative HPLC profiles of the chromophore of ChRmine in the dark (top), under illumination (middle), and after light adaptation (bottom). “at”, “11”, and “13” indicate the peak of all*-trans-*, 11*-cis-*, and 13*-cis-*retinal oximes, respectively. **b**, Calculated composition of all*-trans-* and 13*-cis-*retinal oximes. Data are presented as mean ± s.e.m. (n = 3). Purified samples of ChRmine were prepared with additional supplementation of all*-trans-*retinal. Green light (530 ± 5 nm) was used for illumination. Light adaptation was achieved by illumination for 1 min followed by incubation in the dark for 2 min.

**Extended Data Fig. 7.**
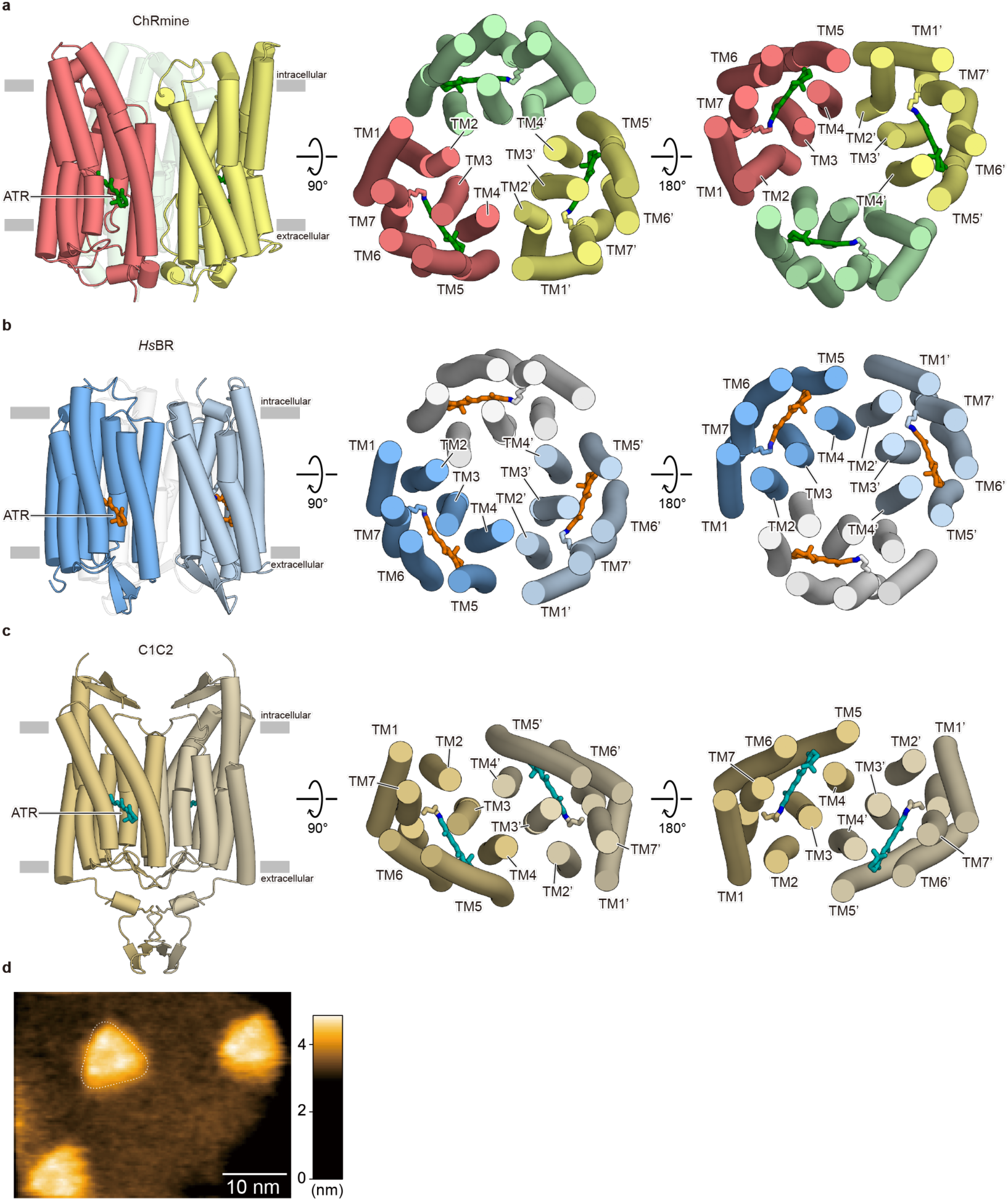
Oligomerization of ChRmine, *Hs*BR, and C1C2. **a**, **b**, **c**, Overall structure of ChRmine (**a**), *Hs*BR (**b**), and C1C2 (**c**), viewed parallel to the membrane (left) and viewed from the intracellular side (middle) and extracellular side (right), shown with each protomer in different colours. **d**, Representative high-speed atomic force microscopy (HS-AFM) image of ChRmine incorporated in the lipid. Frame rate, 2 fps.

**Extended Data Fig. 8.**
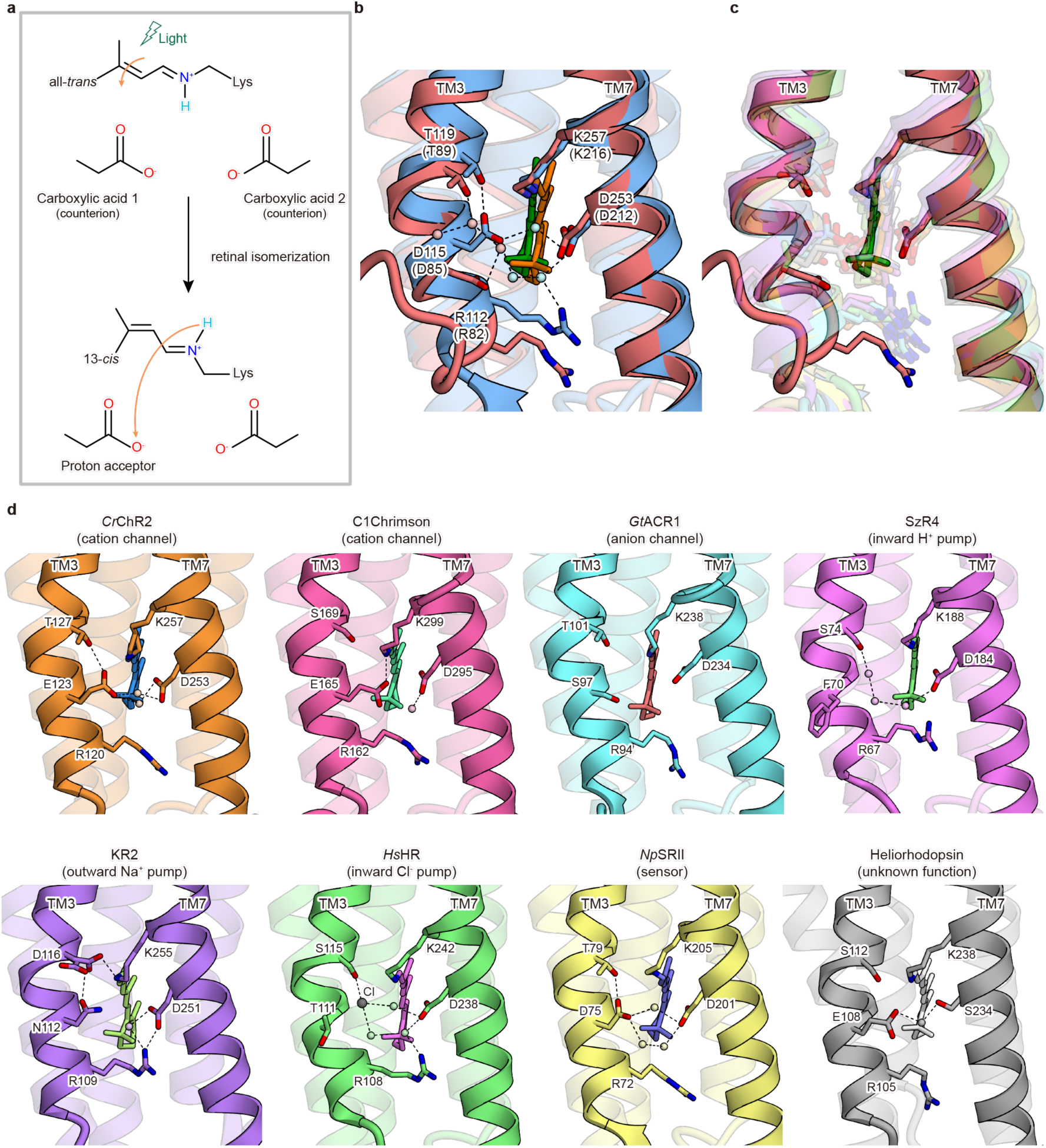
Comparison of the Schiff base regions. **a**, Concept of the Schiff base counterions and proton acceptor. **b**, The superposed Schiff base region of ChRmine (red) and *Hs*BR (blue). Red and blue spheres represent water molecules of ChRmine and *Hs*BR, respectively. Black dashed lines represent hydrogen bonds. **c**, Superposed Schiff base region of ChRmine (red) and representative microbial rhodopsins (*Cr*ChR2, C1Chrimson, *Gt*ACR1, schizorhodopsin 4 (SzR4), KR2, halorhodopsin, *Np*SRII, heliorhodopsin), displayed with high transparency except for ChRmine. **d**, List of the Schiff base regions of representative microbial rhodopsins.

**Extended Data Fig. 9.**
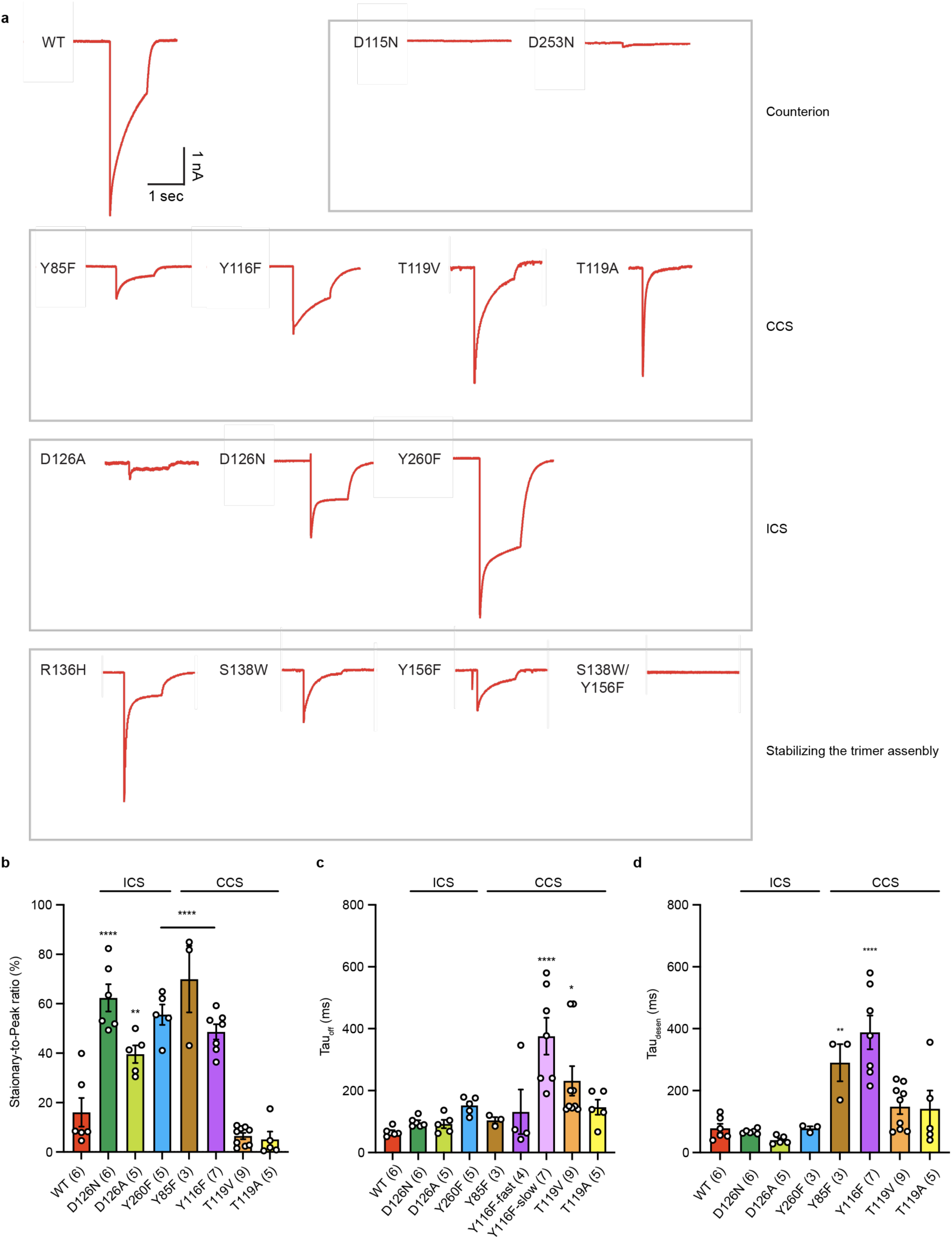
Electrophysiology. **a**, Representative traces of ChRmine WT and 13 mutants expressed in HEK293 cells by lipofectamine transfection, measured at -70 mV holding potential in voltage-clamp. Traces were recorded while cells were stimulated with 1.0 s of 1 mW mm^−2^ irradiance at 580 nm. **b-d**, Summary of the steady-to-peak ratio of photocurrents (**b**), τ_off_ of channel closing (**c**), and τ_off_ of channel desensitization (= τ_desen_) (**d**). Mutants are categorized as the mutants of counterions, central constriction site (CCS), intracellular constriction site (ICS), and the trimer interface. Data are mean ± s.e.m; one-way ANOVA followed by Dunnett’s test. **P < 0.01, ***P < 0.001, and ****P < 0.0001.

**Extended Data Fig. 10.**
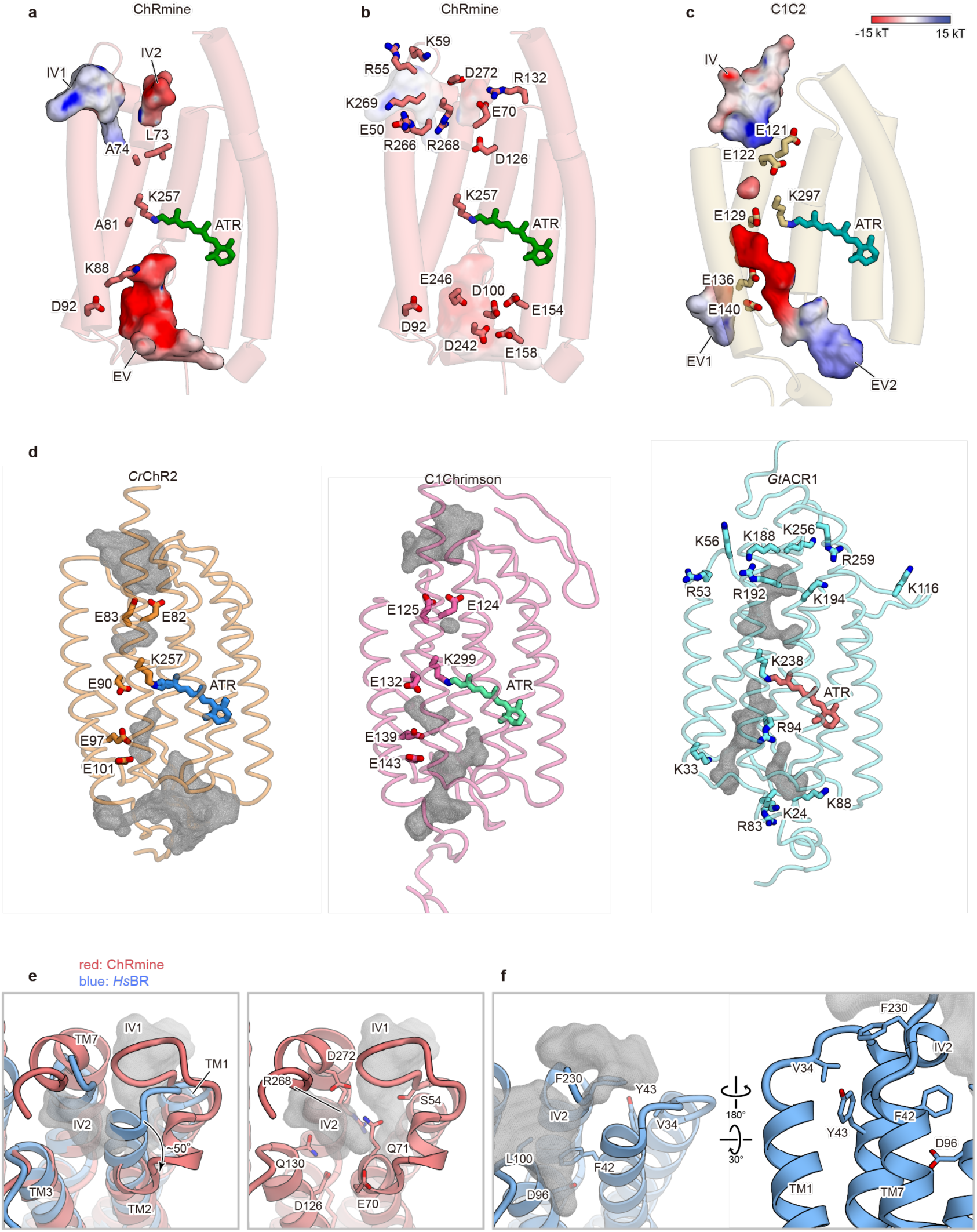
Comparison of ion-conducting pore within the monomer. **a**, **c**, Ion-conducting pore and conserved negatively charged residues (E121, E122, E129, E136, and E140) along the ion-conducting pathway of C1C2 (**c**), and corresponding residues of ChRmine (**a**), as calculated by the software HOLLOW and displayed in Cuemol. Ion-conducting pores are coloured by the electrostatic potential, calculated by PDB2PQR server^74, 75^. **b**, Charged residues along the ion-conducting pore of ChRmine. **d**, Ion conducting pore and conserved negatively charged residues along the ion-conducting pathway of *Cr*ChR2 (left), C1Chrimson (middle), and positively charged residues of *Gt*ACR1 (right). Grey meshes represent the ion-conducting pore. **e**, The superposed intracellular regions of ChRmine (red) and *Hs*BR (blue) (left) and intracellular region of ChRmine (right). The intracellular vestibules (IV1 and IV2) of ChRmine are depicted as grey mesh. **f**, The overall structure of the intracellular region of *Hs*BR. The intracellular vestibules of *Hs*BR are depicted as grey mesh.

**Extended Data Fig. 11.**
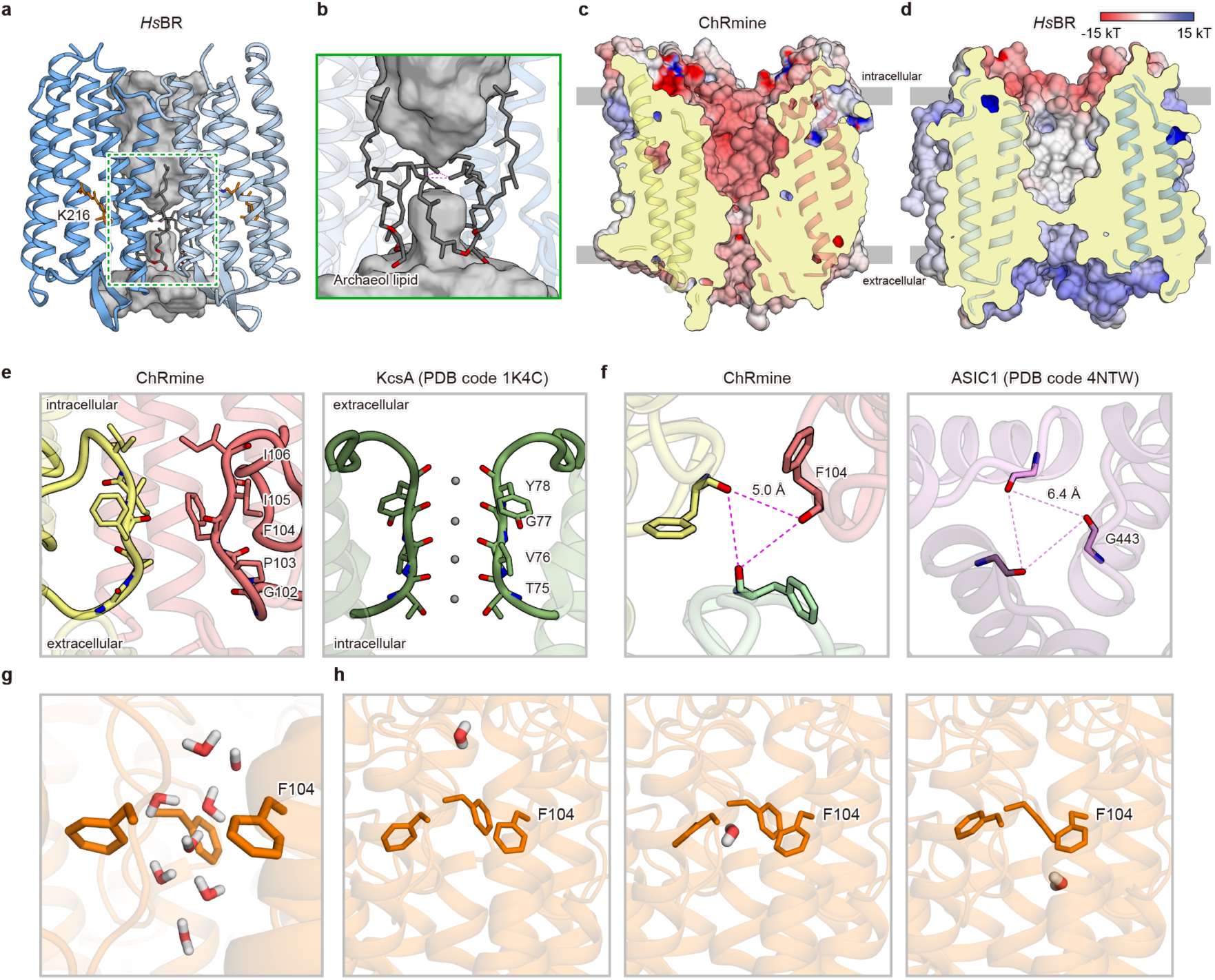
Comparison of ion-conducting pore within the trimer. **a**, Overall structure of the central cavity of the trimer, which is shown as the grey surface. One protomer is hidden for clarity. **b**, Side magnified view of the most constricted site of the central cavity as delimited by the green dashed box. **c**,**d**, Electrostatic potential surface and cross-section of the ChRmine (**c**) and *Hs*BR (**d**). **e,f,** Comparison of ECL of ChRmine with the ion selectivity filter of KcsA (**e**) and ASIC (**f**). **g,h**, Water permeating the trimer interface during the MD simulation. Representative snapshot with water molecules around the constriction (**g**) and successive snapshots focusing on one water molecule (**h**).

**Extended Data Fig. 12.**
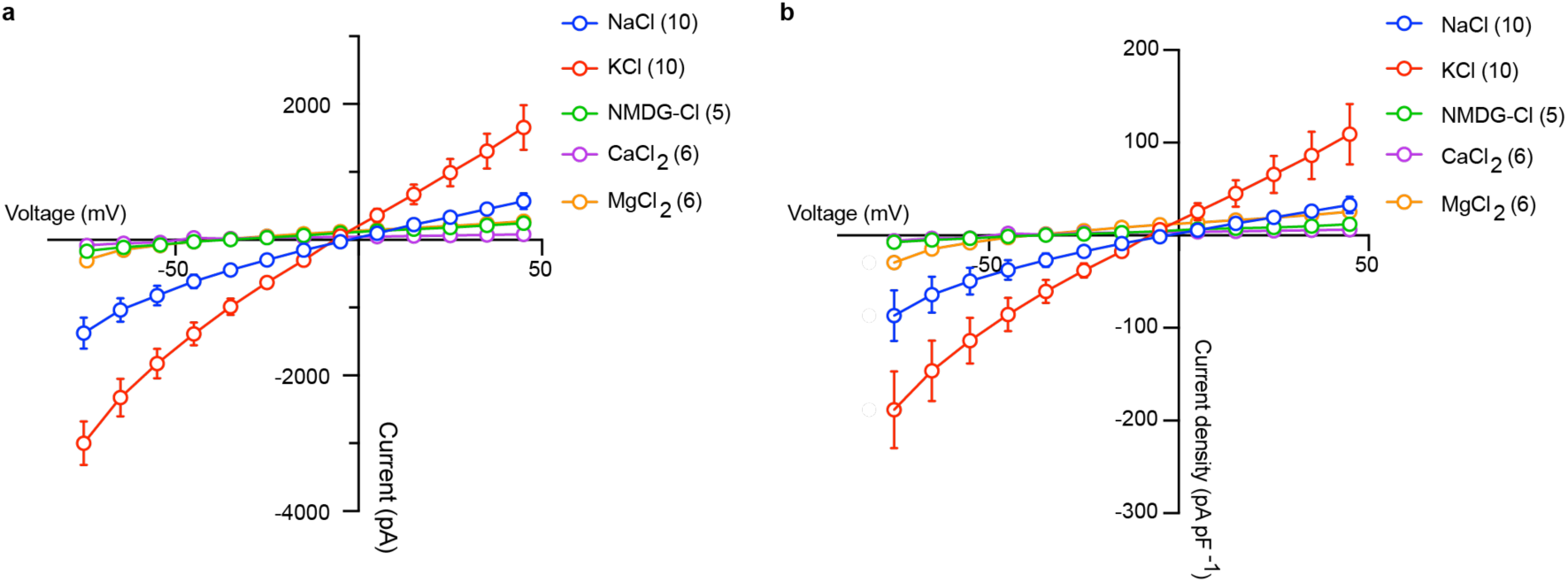
Monovalent cation selectivity of ChRmine. The current-voltage dependence of (**a**) non-normalized and (**b**) normalized peak photocurrents of ChRmine at different voltages from -75 mV to 45 mV stimulated with 1 s of 1 mW mm^−2^ irradiance at 580 nm. Ions in both bath and pipette solutions were replaced with either 120 mM NaCl, 120 mM KCl, 80 mM CaCl_2_, 80 mM MgCl_2_ or 120 mM NMDG-Cl, with all other components at low concentrations (4 mM NaCl, 4 mM KCl, 2 mM CaCl_2_, 2 mM MgCl_2_, and 10 mM HEPES). Data are mean ± s.e.m. Sample size (number of cells) indicated in parentheses.

**Extended Data Fig. 13.**
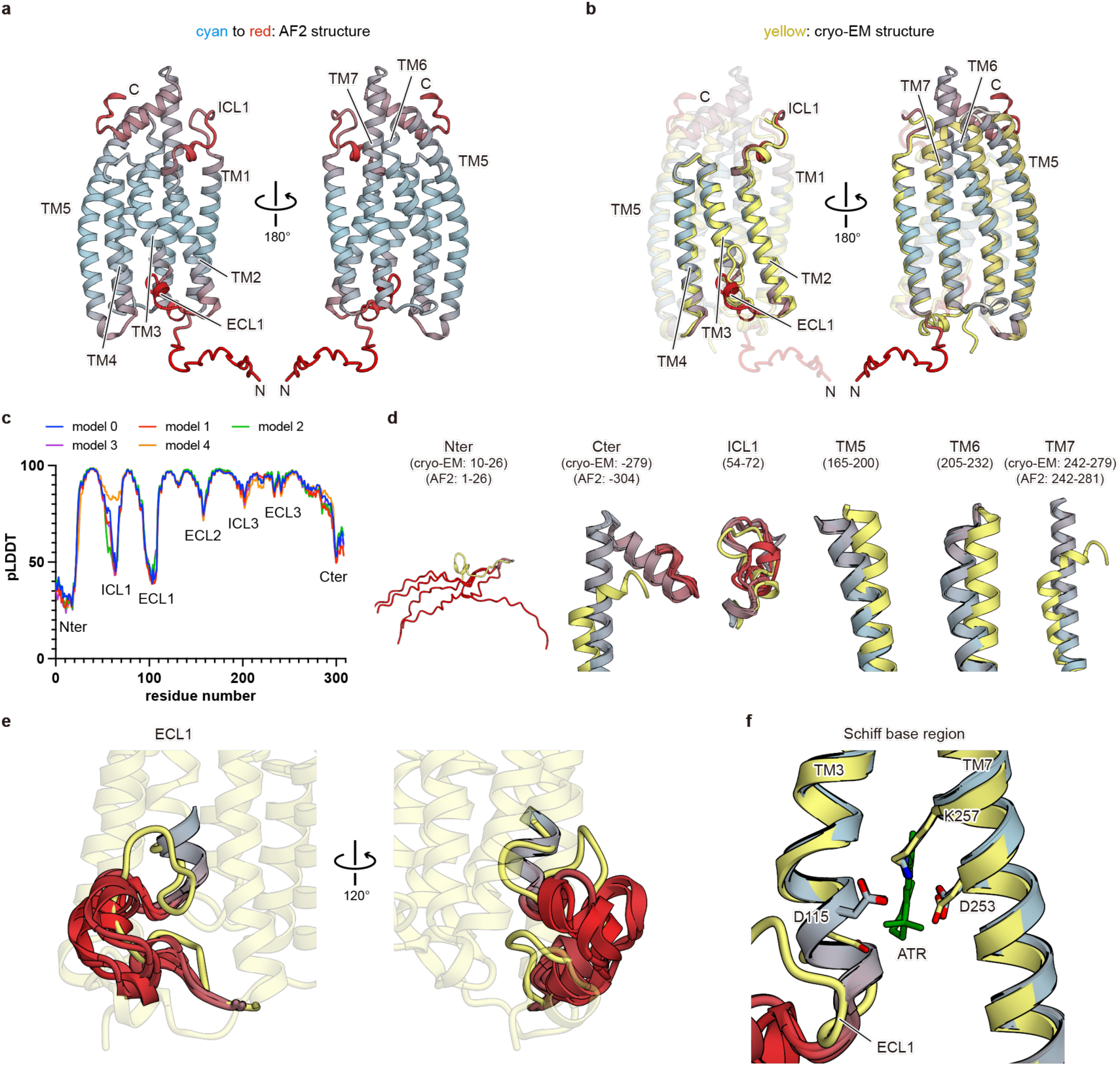
Predicted structures of ChRmine. Five predicted models of ChRmine are generated using locally-installed AlphaFold 2 (https://github.com/deepmind/alphafold). The ribbon representations are coloured by the pLDDT score (low: red, high: cyan). **a**, The best-predicted model of ChRmine, viewed parallel to the membrane. **b**, The predicted model superimposed onto the cryo-EM structure (yellow). **c,** Plots of pLDDT score for five generated models. **d,** The structural comparison of N- and C-terminal regions, ICL1, and TM5-7 between the best predicted model and the cryo-EM structure. Residue numbers are shown in parentheses. **e,f** The detailed comparison of ECL1 (**e**) and the Schiff base region (**f**) between the five predicted models and cryo-EM structure. Notably, the C-terminal region of ECL1, including D115, has high pLDDT scores, but the conformation of D115 is not correctly predicted.

**Extended Data Fig. 14.**
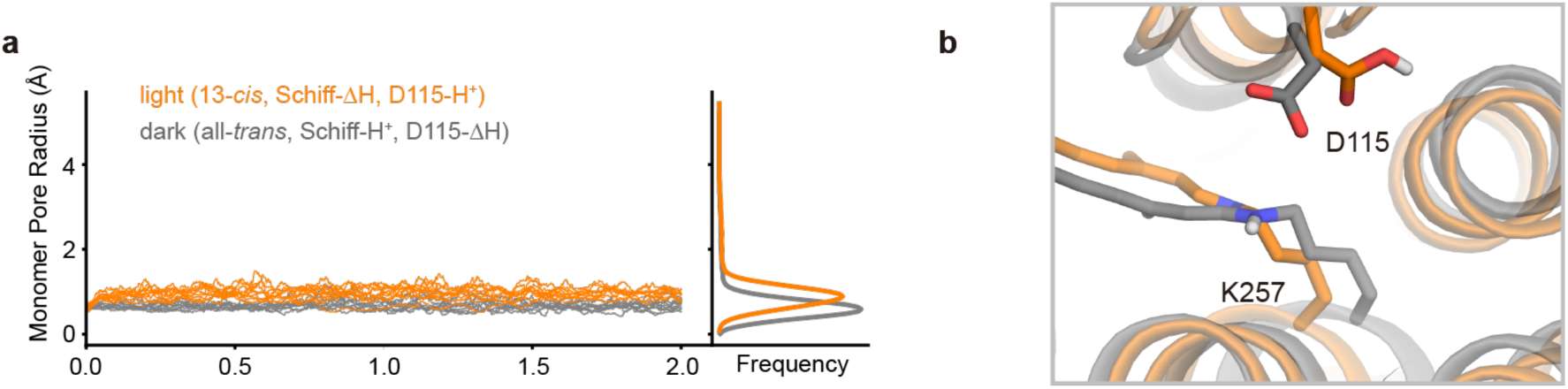
Monomer pore change observed during simulation. **a,** The ChRmine monomer pore opens wider in light-state simulations (13-*cis*- retinal and protonated D115) compared to dark-state simulations (all-*trans*-retinal and deprotonated D115) (p < 0.001, Welch’s t-test). Average minimum monomer pore radius for the three monomers was calculated for each of 10 independent 2 µs simulations. **b**, The pore radius is larger in the light-state simulations than the dark-state simulations as shown through representative frames from the two simulation conditions. Rotation of D115 and isomerization of the retinal opens up the internal monomer space.

**Extended Data Fig. 15.**
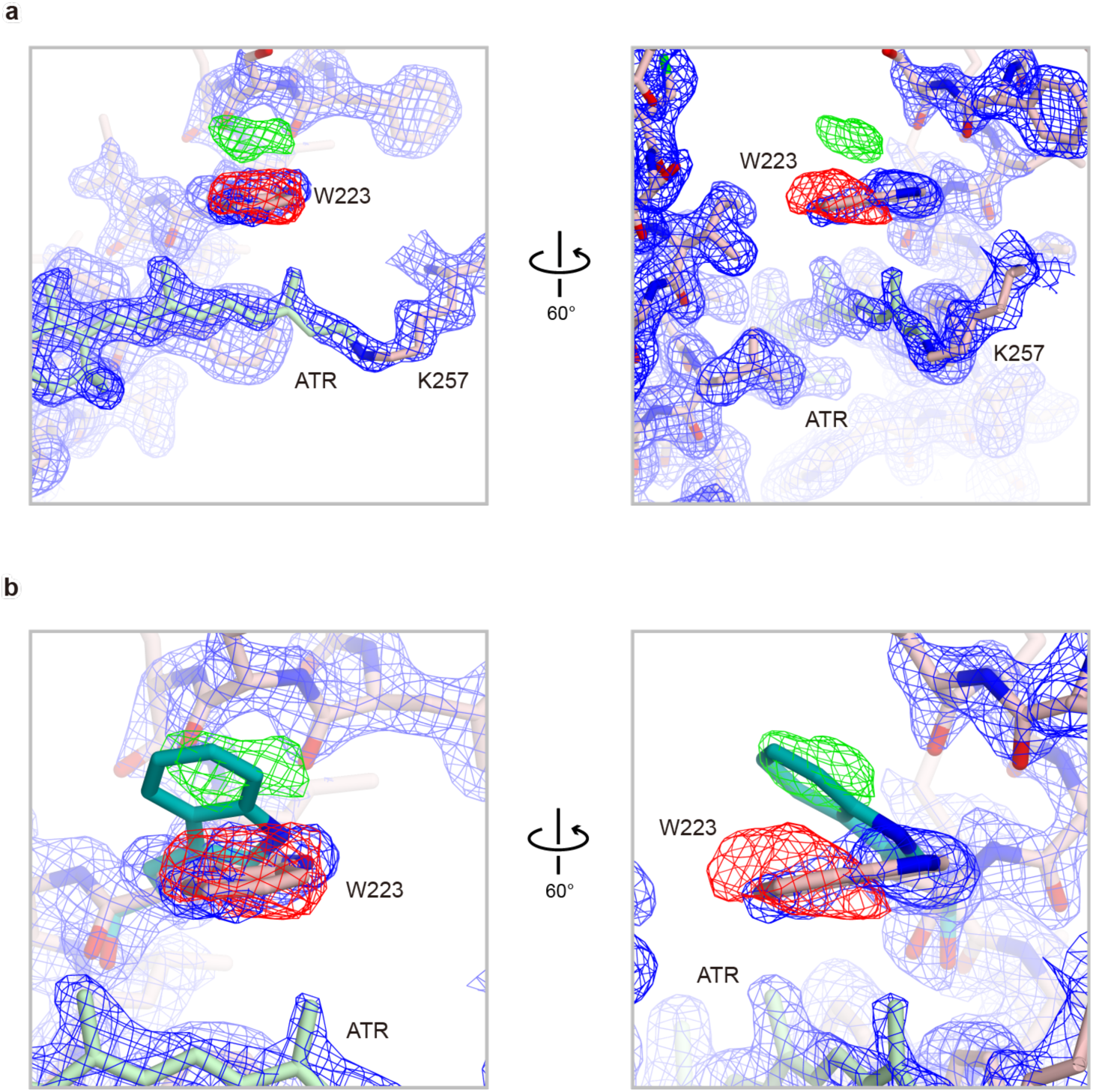
Possible signal of early photo-intermediate. **a**, Density and model in the retinal binding pocket region. Blue and green maps are FSC-weighted sharpened map calculated by RELION3.1.1 and *F_o_-F_c_* maps calculated by Servalcat, respectively. Positive and negative *F_o_-F_c_* difference density pairing (± 3.7σ, where σ is the standard deviation within the mask) is observed around W223, suggesting that this density contains information regarding a small population of the early intermediate state, and that W223 moves upward early in the photocycle. **b**, Magnified views of **a.**

**Supplementary Table 1.** Lifetime of each intermediate in ChRmine WT and mutants.

|  | Dark | (K) | L<sub>1</sub> | M<sub>1</sub>/L<sub>2</sub> | M<sub>2</sub> | Dark |
| --- | --- | --- | --- | --- | --- | --- |
| WT | - | << 500 ns | 5.0 ± 0.4 ms | 28 ± 2 ms | 1.09 ± 0.06 s |  |
| WT-Fab | - | << 500 ns | 5.2 ± 0.5 ms | 31 ± 2 ms | 0.94 ± 0.04 s |  |
| D115N | - | << 500 ns | 5.0 ± 0.3 ms | 12.9 ± 0.7 ms | 190 ± 40 ms |  |
| D253N | - | << 500 ns | 1.8 ± 0.2 ms | 55 ± 1 ms | 1.10 ± 0.04 s |  |

**Supplementary Table 2.**
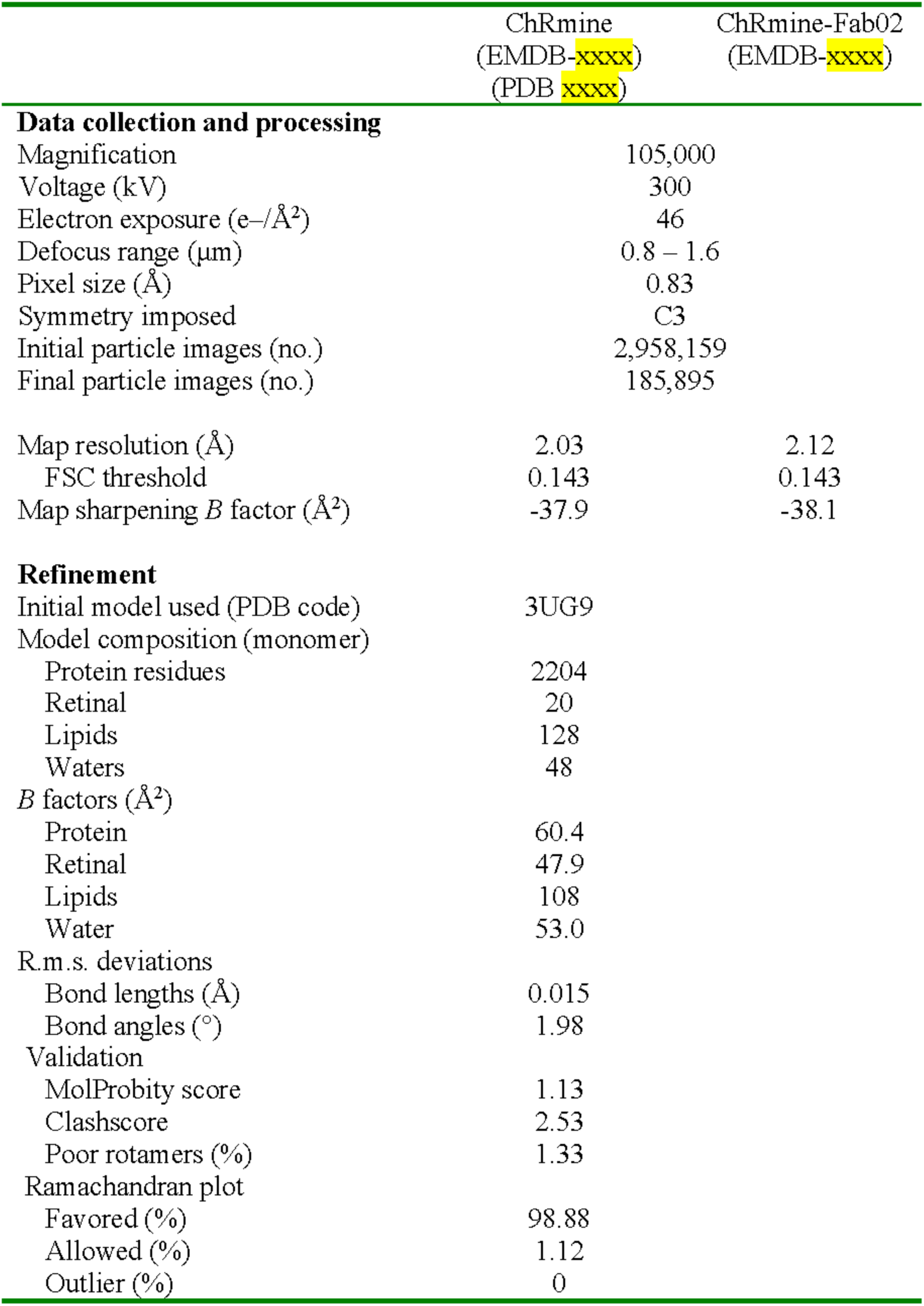
Cryo-EM data collection, refinement, and validation statistics.

## References

1. Marshel, J. H. et al. Cortical layer–specific critical dynamics triggering perception. Science 365, eaaw5202 (2019).

2. Sineshchekov, O. A., Govorunova, E. G., Li, H. & Spudich, J. L. Bacteriorhodopsin-like channelrhodopsins: Alternative mechanism for control of cation conductance. Proc. Natl. Acad. Sci. 114, E9512–E9519 (2017).

3. Chen, R. et al. Deep brain optogenetics without intracranial surgery. Nat. Biotechnol. 39, 161–164 (2021).

4. Matsubara, T. et al. Remote control of neural function by X-ray-induced scintillation. Nat. Commun. 12, (2021).

5. Okada, T. et al. Pain induces stable, active microcircuits in the somatosensory cortex that provide a therapeutic target. Sci. Adv. 7, eabd8261 (2021).

6. Ernst, O. P. et al. Microbial and animal rhodopsins: Structures, functions, and molecular mechanisms. Chem. Rev. 114, 126–163 (2014).

7. Kandori, H. Biophysics of rhodopsins and optogenetics. Biophys. Rev. 12, 355– 361 (2020).

8. Kato, H. E. Structure-Function Relationship of Channelrhodopsins. Adv. Exp. Med. Biol. 1293, 35–53 (2021).

9. Deisseroth, K. Optogenetics: 10 years of microbial opsins in neuroscience. Nat. Neurosci. 18, 1213–25 (2015).

10. Kurihara, M. & Sudo, Y. Microbial rhodopsins: wide distribution, rich diversity and great potential. Biophys. Physicobiology 12, 121–129 (2015).

11. Deisseroth, K. & Hegemann, P. The form and function of channelrhodopsin. Science 357, eaan5544 (2017).

12. Nagel, G. et al. Channelrhodopsin-1: A light-gated proton channel in green algae. Science 296, 2395–2398 (2002).

13. Nagel, G. et al. Channelrhodopsin-2, a directly light-gated cation-selective membrane channel. Proc. Natl. Acad. Sci. 100, 13940–13945 (2003).

14. Klapoetke, N. C. et al. Independent optical excitation of distinct neural populations. Nat. Methods 11, 338–346 (2014).

15. Govorunova, E. G., Sineshchekov, O. A. & Spudich, J. L. Structurally Distinct Cation Channelrhodopsins from Cryptophyte Algae. Biophys. J. 110, 2302–2304 (2016).

16. Yamauchi, Y. et al. Molecular properties of a DTD channelrhodopsin from *Guillardia theta*. Biophys. Physicobiology 14, 57–66 (2017).

17. Inoue, K. et al. A light-driven sodium ion pump in marine bacteria. Nat. Commun. 4, 1–10 (2013).

18. Kato, H. E. et al. Crystal structure of the channelrhodopsin light-gated cation channel. Nature 482, 369–374 (2012).

19. Berndt, A., Lee, S. Y., Ramakrishnan, C. & Deisseroth, K. Structure-Guided Transformation of Channelrhodopsin into a Light-Activated Chloride Channel. Science 344, 420–424 (2014).

20. Wietek, J. et al. Conversion of Channelrhodopsin into a Light-Gated Chloride Channel. Science 344, 409–412 (2014).

21. Berndt, A. et al. Structural foundations of optogenetics: Determinants of channelrhodopsin ion selectivity. Proc. Natl. Acad. Sci. 113, 822–829 (2016).

22. Yamashita, K., Palmer, C. M., Burnley, T. & Murshudov, G. N. Structural Biology Cryo-EM single particle structure refinement and map calculation using Servalcat. bioRxiv 2021.05.04.442493 (2021). doi:10.1101/2021.05.04.442493

23. Pebay-Peyroula, E., Rummel, G., Rosenbusch, J. P. & Landau, E. M. X-ray structure of bacteriorhodopsin at 2.5 angstroms from microcrystals grown in lipidic cubic phases. Science 277, 1676–81 (1997).

24. Kolbe, M., Besir, H., Essen, L. O. & Oesterhelt, D. Structure of the light-driven chloride pump halorhodopsin at 1.8 A resolution. Science 288, 1390–6 (2000).

25. Gerwert, K., Souvignier, G. & Hess, B. Simultaneous monitoring of light-induced changes in protein side-group protonation, chromophore isomerization, and backbone motion of bacteriorhodopsin by time-resolved Fourier-transform infrared spectroscopy. Proc. Natl. Acad. Sci. 87, 9774–9778 (1990).

26. Lorenz-Fonfria, V. A. et al. Transient protonation changes in channelrhodopsin-2 and their relevance to channel gating. Proc. Natl. Acad. Sci. 110, E1273–E1281 (2013).

27. Gunaydin, L. A. et al. Ultrafast optogenetic control. Nat. Neurosci. 13, 387–392 (2010).

28. Berndt, A. & Deisseroth, K. Expanding the optogenetics toolkit. Science 349, 590– 591 (2015).

29. Takemoto, M. et al. Molecular dynamics of channelrhodopsin at the early stages of channel opening. PLoS One 10, 1–15 (2015).

30. Kato, H. E. et al. Structural mechanisms of selectivity and gating in anion channelrhodopsins. Nature 561, 349–354 (2018).

31. Fudim, R. et al. Design of a light-gated proton channel based on the crystal structure of Coccomyxa rhodopsin. Sci. Signal. 12, eaav4203 (2019).

32. Vogt, A. et al. Engineered Passive Potassium Conductance in the KR2 Sodium Pump. Biophys. J. 116, 1941–1951 (2019).

33. Kawate, T. & Gouaux, E. Fluorescence-Detection Size-Exclusion Chromatography for Precrystallization Screening of Integral Membrane Proteins. Structure 14, 673–681 (2006).

34. Sineshchekov, O. A. et al. Conductance mechanisms of rapidly desensitizing cation channelrhodopsins from cryptophyte algae. MBio 11, 1–12 (2020).

35. Jumper, J. et al. Highly accurate protein structure prediction with AlphaFold. Nature (2021). doi:10.1038/s41586-021-03819-2

36. Mariani, V., Biasini, M., Barbato, A. & Schwede, T. lDDT: a local superposition-free score for comparing protein structures and models using distance difference tests. Bioinformatics 29, 2722–2728 (2013).

37. Tunyasuvunakool, K. et al. Highly accurate protein structure prediction for the human proteome. Nature (2021). doi:10.1038/s41586-021-03828-1

## Additional References

38. Sievers, F. & Higgins, D. G. Clustal Omega. Curr. Protoc. Bioinforma. 48, 3.13.1-3.13.16 (2014).

39. Zivanov, J. et al. New tools for automated high-resolution cryo-EM structure determination in RELION-3. Elife 7, (2018).

40. Zheng, S. Q. et al. MotionCor2: Anisotropic correction of beam-induced motion for improved cryo-electron microscopy. Nature Methods 14, 331–332 (2017).

41. Zivanov, J., Nakane, T. & Scheres, S. H. W. A Bayesian approach to beam-induced motion correction in cryo-EM single-particle analysis. IUCrJ 6, 5–17 (2019).

42. Zivanov, J., Nakane, T. & Scheres, S. H. W. Estimation of high-order aberrations and anisotropic magnification from cryo-EM data sets in RELION-3.1. IUCrJ 7, 253–267 (2020).

43. Ramlaul, K., Palmer, C. M., Nakane, T. & Aylett, C. H. S. Mitigating local over-fitting during single particle reconstruction with SIDESPLITTER. J. Struct. Biol. 211, 107545 (2020).

44. Emsley, P. & Cowtan, K. Coot : model-building tools for molecular graphics. Acta Crystallogr. Sect. D Biol. Crystallogr. 60, 2126–2132 (2004).

45. Murshudov, G. N. et al. REFMAC5 for the refinement of macromolecular crystal structures. Acta Crystallogr. Sect. D Biol. Crystallogr. 67, 355–367 (2011).

46. Chen, V. B. et al. MolProbity: All-atom structure validation for macromolecular crystallography. Acta Crystallogr. Sect. D Biol. Crystallogr. 66, 12–21 (2010).

47. Pettersen, E. F. et al. UCSF Chimera - A visualization system for exploratory research and analysis. J. Comput. Chem. 25, 1605–1612 (2004).

48. Goddard, T. D. et al. UCSF ChimeraX: Meeting modern challenges in visualization and analysis. Protein Sci. 27, 14–25 (2018).

49. Trehan, A. et al. On retention of chromophore configuration of rhodopsin isomers derived from three dicis retinal isomers. Bioorg. Chem. 18, 30–40 (1990).

50. Bayburt, T. H., Grinkova, Y. V. & Sligar, S. G. Self-Assembly of Discoidal Phospholipid Bilayer Nanoparticles with Membrane Scaffold Proteins. Nano Lett. 2, 853–856 (2002).

51. Denisov, I. G. & Sligar, S. G. Nanodiscs for structural and functional studies of membrane proteins. Nat. Struct. Mol. Biol. 23, 481–486 (2016).

52. Shibata, M. et al. Oligomeric states of microbial rhodopsins determined by high-speed atomic force microscopy and circular dichroic spectroscopy. Sci. Rep. 8, 1– 11 (2018).

53. Shibata, M. et al. Real-space and real-Time dynamics of CRISPR-Cas9 visualized by high-speed atomic force microscopy. Nat. Commun. 8, 1–9 (2017).

54. Betz, R. Dabble. (2017). doi:10.5281/zenodo.836914.

55. Hopkins, C. W., Le Grand, S., Walker, R. C. & Roitberg, A. E. Long-Time-Step Molecular Dynamics through Hydrogen Mass Repartitioning. J. Chem. Theory Comput. 11, 1864–1874 (2015).

56. Ryckaert, J.-P., Ciccotti, G. & Berendsen, H. J. C. Numerical integration of the cartesian equations of motion of a system with constraints: molecular dynamics of n-alkanes. J. Comput. Phys. 23, 327–341 (1977).

57. Beglov, D. & Roux, B. Finite representation of an infinite bulk system: Solvent boundary potential for computer simulations. J. Chem. Phys. 100, 9050–9063 (1994).

58. Huang, J. et al. CHARMM36m: an improved force field for folded and intrinsically disordered proteins. Nat. Methods 14, 71–73 (2017).

59. Klauda, J. B. et al. Update of the CHARMM All-Atom Additive Force Field for Lipids: Validation on Six Lipid Types. J. Phys. Chem. B 114, 7830–7843 (2010).

60. Roe, D. R. & Cheatham, T. E. PTRAJ and CPPTRAJ: Software for Processing and Analysis of Molecular Dynamics Trajectory Data. J. Chem. Theory Comput. 9, 3084–3095 (2013).

61. Humphrey, W., Dalke, A. & Schulten, K. VMD: Visual molecular dynamics. J. Mol. Graph. 14, 33–38 (1996).

62. GitHub - osmart/hole2: Source code for HOLE program. Available at: https://github.com/osmart/hole2.

63. Kim, Y. S. et al. Crystal structure of the natural anion-conducting channelrhodopsin GtACR1. Nature 561, 343–348 (2018).

64. Hasegawa, N., Jonotsuka, H., Miki, K. & Takeda, K. X-ray structure analysis of bacteriorhodopsin at 1.3 Å resolution. Sci. Rep. 8, 1–8 (2018).

65. Volkov, O. et al. Structural insights into ion conduction by channelrhodopsin 2. Science 358, eaan8862 (2017).

66. Oda, K. et al. Crystal structure of the red light-activated channelrhodopsin Chrimson. Nat. Commun. 9, 1–11 (2018).

67. Ran, T. et al. Cross-protomer interaction with the photoactive site in oligomeric proteorhodopsin complexes. Acta Crystallogr. Sect. D Biol. Crystallogr. 69, 1965– 1980 (2013).

68. Luecke, H. et al. Crystallographic structure of xanthorhodopsin, the light-driven proton pump with a dual chromophore. Proc. Natl. Acad. Sci. U. S. A. 105, 16561– 16565 (2008).

69. Kato, H. E. et al. Structural basis for Na(+) transport mechanism by a light-driven Na(+) pump. Nature 521, 48–53 (2015).

70. Pei, J., Kim, B. H. & Grishin, N. V. PROMALS3D: A tool for multiple protein sequence and structure alignments. Nucleic Acids Res. 36, 2295–2300 (2008).

71. Robert, X. & Gouet, P. Deciphering key features in protein structures with the new ENDscript server. Nucleic Acids Res. 42, W320–W324 (2014).

72. Saitou, N. & Nei, M. The neighbor-joining method: a new method for reconstructing phylogenetic trees. Mol. Biol. Evol. 4, 406–425 (1987).

73. Kumar, S., Stecher, G. & Tamura, K. MEGA7: Molecular Evolutionary Genetics Analysis Version 7.0 for Bigger Datasets. Mol. Biol. Evol. 33, 1870–1874 (2016).

74. Baker, N. A., Sept, D., Joseph, S., Holst, M. J. & McCammon, J. A. Electrostatics of nanosystems: Application to microtubules and the ribosome. Proc. Natl. Acad. Sci. 98, 10037–10041 (2001).

75. Dolinsky, T. J., Nielsen, J. E., McCammon, J. A. & Baker, N. A. PDB2PQR: an automated pipeline for the setup of Poisson-Boltzmann electrostatics calculations. Nucleic Acids Res. 32, W665–W667 (2004).

