## Supplementary Information for "Structural basis for channel conduction in the pump-like channelrhodopsin ChRmine"

**Supplementary Discussion**

**Arginine conformation in the dark state and the function of ion-transporting rhodopsins.**

As shown here and in previous studies, R82 in *Hs*BR is highly conserved among microbial rhodopsins, but two different conformations are seen in the dark state: outward-facing and parallel. In all structurally-resolved channelrhodopsins (C1C2 (PDB ID: 3UG9)[1], *Cr*ChR2 (PDB ID: 6EID)[2], C1Chrimson (PDB ID: 5ZIH)[3], *Gt*ACR1 (PDB ID: 6CSM)[4], and ChRmine), the arginine residue faces outward. In contrast, in many ion-pumping rhodopsins, including *Hs*BR (PDB ID: 5ZIM)[5], *Hw*BR (PDB ID: 4QID), cruxrhodopsin-3 (PDB ID: 4JR8)[6], deltarhodopsin (PDB ID: 4FBZ)[7], GR (PDB ID: 6NWD)[8], Archaerhodopsin-1 (PDB ID: 1UAZ)[9], Archaerhodopsin-2 (PDB ID: 2EI4)[10], PR from the Mediterranean Sea at a depth of 12 m (*Med12*BPR, PDB ID: 4JQ6)[11], PR from the Pacific Ocean near Hawaii at a depth of 75 m (*HOT75*BPR, PDB ID: 4KLY)[11], *Cs*R (6GYH)[12], *Hs*HR (PDB ID: 1E12)[13], *Np*HR (PDB ID: 3A7K)[14], ClR (PDB ID: 5ZTK)[15], KR2 (PDB ID: 3X3C)[16], and schizorhodopsin 4 (PDB ID: 7E4G)[17], the tip of arginine runs parallel to the membrane and faces TM1. The arginine in the parallel conformation narrows or blocks the extracellular cavity of the ion-translocating pathway; thus, this conformation would contribute to preventing large ion flux in ion-pumping rhodopsins.

Notably, *Cs*R (the outward proton-pumping rhodopsin from *Coccomyxa subellipsoidea*), also has arginine (R83) in the parallel conformation in the dark state[12], and R83Q mutation or mutation of the adjacent tyrosine (Y57K) converts functionality from proton pump to proton channel[18]. Moreover, computational analysis of *Hs*BR with R82Q or Y57K mutation reveals that these mutations significantly change the conformation of R82Q or R82, respectively; most notably, R82 faces outward in the Y57K simulation[18]. These observations suggest that the conformation of the arginine in the dark state is one of the structural elements distinguishing channel- and pump-type rhodopsins. Interestingly, previous studies have reported that the arginine of some ion-pumping rhodopsins is maintained in the parallel conformation during the photocycle[19,20], but the corresponding arginine in *Hs*BR transiently changes from parallel to outward-facing to facilitate proton release to the extracellular solvent[21,22]. Since channelrhodopsins presumably evolved from ion-pumping rhodopsins[23], these studies suggest that mutations accumulated near the arginine of ion-pumping rhodopsins gradually stabilized the outward-facing conformation; these rearrangements enlarged the extracellular cavity, enabling the large ion flux of channelrhodopsins.

**The conformational change of the monomer pore during simulation.**

In addition to the opening of the trimer pore, we also observed that the size of the monomer pore increased during the light state simulation (Extended Data Fig. 14a). Upon isomerization and proton transfer, both the retinal and D115 rotate away from the internal monomer pore, increasing the space within the monomer (Extended Data Fig. 14b). While over the timescale of our simulation, the pore radius does not become large enough to allow for travel of ions through the monomers (as with the trimer pore), the monomeric changes follow the expected opening that would occur upon light activation.

**Proton donor and acceptor.**

In *Hs*BR, D85 receives a proton from the protonated Schiff base and releases it to the extracellular bulk solvent. D96 receives a proton from the intracellular bulk solvent and provides it to the deprotonated Schiff base. These proton movements generate net flow of proton from the intracellular to extracellular side; these two functionally important residues, together with T89, are called DTD motif.

In *Gt*CCR2, a ChRmine homolog in the BCCR family, both D85 and D96 are conserved (D87 and D98, respectively) but the proposed proton translocation pathway is completely different; *Gt*CCR2 does not show outward proton-pumping activity[24], and the proton is shuttled back and forth between the Schiff base and D85. While the deprotonation and re-protonation of D98 are assumed to occur and the deprotonation would be necessary for the channel gating, D98 never gives the proton to the deprotonated Schiff base[24].

If some channelrhodopsins retain residual pumping activity[25], D85 homologs presumably could release a proton to the extracellular bulk solvent after receiving a proton from the Schiff base. However, in contrast to the proton acceptor, it remains elusive which residue works as a proton donor to the deprotonated Schiff base. D96 in *Hs*BR is also conserved in ChRmine (D126), but is exposed to intracellular bulk solvent in our structure, and the calculated pKa of D126 is as low as 6.28. D126 may be unlikely to work as the sole proton donor (Fig. 3c) although is possible that the deprotonated Schiff base directly receives a proton from water molecules. Future studies will be needed to reveal the molecular nature of the proton donor, and understand how proton translocation is involved in channel gating of ChRmine.

**A small population of early intermediates: initial structural changes in ChRmine.**

Our 2.0 Å cryo-EM map allowed accurate modeling of ATR and surrounding residues, but the C13 and C14 atoms of ATR and W223 showed weaker density in the region. Moreover, positive and negative $F_o$-$F_c$ difference densities were observed around W223, suggesting that this cryo-EM density map contains information on a small population of early intermediate states (possibly the K intermediate; (Extended Data Fig. 15). While we could not detect further structural changes propagated from W223, the extent of the conformational change in ChRmine W223 was significantly larger than that in C1C2 and similar to that of *Hs*BR[26,27]. Thus, it is concluded that the initial conformational changes of the ChRmine photocycle may be more similar to those of ion-pumping rhodopsins than of canonical chlorophyte channelrhodopsins.

### References


1. Kato, H. E. *et al.* Crystal structure of the channelrhodopsin light-gated cation channel. *Nature* **482**, 369–374 (2012).
2. Volkov, O. *et al.* Structural insights into ion conduction by channelrhodopsin 2. *Science.* **358**, eaan8862 (2017).
3. Oda, K. *et al.* Crystal structure of the red light-activated channelrhodopsin Chrimson. *Nat. Commun.* **9**, 1–11 (2018).
4. Kim, Y. S. *et al.* Crystal structure of the natural anion-conducting channelrhodopsin *Gt*ACR1. *Nature* **561**, 343–348 (2018).
5. Hasegawa, N., Jonotsuka, H., Miki, K. & Takeda, K. X-ray structure analysis of bacteriorhodopsin at 1.3 Å resolution. *Sci. Rep.* **8**, 1–8 (2018).
6. Chan, S. K. *et al.* Crystal structure of cruxrhodopsin-3 from haloarcula vallismortis. *PLoS One* **9**, (2014).
7. Zhang, J. *et al.* Crystal structure of deltarhodopsin-3 from Haloterrigena thermotolerans. *Proteins Struct. Funct. Bioinforma.* **81**, 1585–1592 (2013).
8. Morizumi, T. *et al.* X-ray Crystallographic Structure and Oligomerization of Gloeobacter Rhodopsin. *Sci. Rep.* **9**, 11283 (2019).
9. Enami, N. *et al.* Crystal Structures of Archaerhodopsin-1 and -2: Common Structural Motif in Archaeal Light-driven Proton Pumps. *J. Mol. Biol.* **358**, 675–685 (2006).
10. Yoshimura, K. & Kouyama, T. Structural Role of Bacterioruberin in the Trimeric Structure of Archaerhodopsin-2. *J. Mol. Biol.* **375**, 1267–1281 (2008).
11. Ran, T. *et al.* Cross-protomer interaction with the photoactive site in oligomeric proteorhodopsin complexes. *Acta Crystallogr. Sect. D Biol. Crystallogr.* **69**, 1965–1980 (2013).
12. Fudim, R. *et al.* Design of a light-gated proton channel based on the crystal structure of Coccomyxa rhodopsin. *Sci. Signal.* **12**, eaav4203 (2019).
13. Kolbe, M. Structure of the Light-Driven Chloride Pump Halorhodopsin at 1.8 Å Resolution. *Science.* **288**, 1390–1396 (2000).
14. Kouyama, T. *et al.* Crystal Structure of the Light-Driven Chloride Pump Halorhodopsin from Natronomonas pharaonis. *J. Mol. Biol.* **396**, 564–579 (2010).
15. Yun, J.-H. *et al.* Early-stage dynamics of chloride ion–pumping rhodopsin revealed by a femtosecond X-ray laser. *Proc. Natl. Acad. Sci.* **118**, e2020486118 (2021).
16. Kato, H. E. *et al.* Structural basis for Na(+) transport mechanism by a light-driven Na(+) pump. *Nature* **521**, 48–53 (2015).
17. Higuchi, A. *et al.* Crystal structure of schizorhodopsin reveals mechanism of inward proton pumping. *Proc. Natl. Acad. Sci. U. S. A.* **118**, 1–10 (2021).

18. Vogt, A. *et al.* Conversion of a light-driven proton pump into a light-gated ion channel. *Sci. Rep.* **5**, 16450 (2015).
19. Kouyama, T., Kawaguchi, H., Nakanishi, T., Kubo, H. & Murakami, M. Crystal Structures of the L 1 , L 2 , N, and O States of pharaonis Halorhodopsin. *Biophys. J.* **108**, 2680–2690 (2015).
20. Kovalev, K. *et al.* Molecular mechanism of light-driven sodium pumping. *Nat. Commun.* **11**, 2137 (2020).
21. Kühlbrandt, W. Bacteriorhodopsin — the movie. *Nature* **406**, 569–570 (2000).
22. Nango, E. *et al.* A three-dimensional movie of structural changes in bacteriorhodopsin. *Science.* **354**, 1552–1557 (2016).
23. Inoue, K. *et al.* Converting a Light-Driven Proton Pump into a Light-Gated Proton Channel. *J. Am. Chem. Soc.* **137**, 3291–3299 (2015).
24. Sineshchekov, O. A., Govorunova, E. G., Li, H. & Spudich, J. L. Bacteriorhodopsin-like channelrhodopsins: Alternative mechanism for control of cation conductance. *Proc. Natl. Acad. Sci.* **114**, E9512–E9519 (2017).
25. Feldbauer, K. *et al.* Channelrhodopsin-2 is a leaky proton pump. *Proc. Natl. Acad. Sci.* **106**, 12317–12322 (2009).
26. Weinert, T. *et al.* Proton uptake mechanism in bacteriorhodopsin captured by serial synchrotron crystallography. *Science.* **365**, 61–65 (2019).
27. Oda, K. *et al.* Time-resolved serial femtosecond crystallography reveals early structural changes in channelrhodopsin. *Elife* **10**, (2021).
